# Cassette recruitment in the chromosomal Integron of *Vibrio cholerae*

**DOI:** 10.1101/2020.11.24.395434

**Authors:** Claire Vit, Egill Richard, Florian Fournes, Clémence Whiteway, Xavier Eyer, Delphine Lapaillerie, Vincent Parissi, Didier Mazel, Céline Loot

## Abstract

Integrons are genetic systems conferring to bacteria a rapid adaptation capability. The integron integrase is able to capture, stockpile and shuffle novel functions embedded in cassettes. This involves the recognition of both substrates, the *attI* site, and the cassette associated *attC* sites. Integrons can be sedentary and chromosomally located (SCI) or, carried by conjugative plasmids (Mobile Integron, MI), hence favoring their dissemination among bacteria. Here, for the first time, we investigate the cassette recruitment in the *Vibrio cholerae* SCI during conjugation and natural transformation. We demonstrated that horizontally transferred cassette can be recruited inside the chromosomal integron. The endogenous integrase expression is sufficiently triggered, after SOS response induction mediated by the entry of single-stranded cassettes during conjugation and natural transformation, to mediate significant cassette insertion. We demonstrate that the *attIA* insertion is preferential, despite the presence of 180 *attC* sites in the integron array. Thanks to the presence of a promoter in the *attIA* site vicinity, all these newly inserted cassettes are expressed and prone to adaptive selection. We also show that the RecA protein is critical for cassette recruitment in *V. cholerae* SCI but not in MIs. Moreover, *a contrario* to MIs, the *V. cholerae* SCI is not active in others bacterial hosts. MIs might have evolved from the SCIs by overcoming host factors, which would explain their large dissemination in bacteria and their role in the antibioresistance expansion.

## INTRODUCTION

Mobile Genetic Elements (MGE) widely contribute to the evolution of bacterial genomes notably by conferring adaptive traits such as the ability to resist antibiotic treatments (Partridge et al., 2018). This can have dramatic consequences, especially when concerning pathogenic bacteria. Integrons are MGE that are considered as major contributors in the rise of multi-resistance in Gram-negative bacteria. These genetic systems were discovered in the late 80s and described as platforms involved in the capture, stockpiling and expression of antibiotic resistance genes, embedded in structures termed “cassettes” (Stokes and Hall, 1989). These latter were referred to as mobile integrons (MIs) because of their associations with transposable elements and conjugative plasmids. Larger integrons located on bacterial chromosomes were discovered later, the superintegron of *V. cholerae* being the first identified (Mazel et al., 1998). This superintegron is located on the chromosome 2 of *V. cholerae* and contains 180 gene cassettes coding mainly for proteins with no homologs in the databases or for proteins of unknown function. In contrast to their mobile counterpart and to refer at their location, such structures are termed Sedentary Chromosomal Integrons (SCI). They are common features of bacterial genomes from *Vibrio* genus and are generally distributed in several genomes of β- and γ-proteobacteria (Cambray et al., 2010; Cury et al., 2016). These large SCIs were proposed to be the ancestors of MIs (Rowe-Magnus et al., 2001) and the stockpiling capacity from both types of integrons suggests that they may have distinct but complementary roles (Loot et al., 2017). Indeed, large SCIs such as those found in genomes of *Vibrio* species, could constitute a reservoir of gene cassettes that can be captured and spread by MIs (Loot et al., 2017; Rowe-Magnus et al., 2002).

Both types of integrons are composed of a stable platform and a variable cassette array. The platform contains a gene coding for a tyrosine-recombinase, IntI, the *attI* recombination site and the resident promoter P_C_ oriented toward the variable cassette array (Figure 1A). The cassette array contains a pool of gene cassettes generally composed of single promoter-less genes (coding sequence, CDS), so they rely on P_C_ promoter to drive their expression. Starting from P_C_, their expression gradually decreases in the array (Jacquier et al., 2009) (Figure 1A). Then, the gene cassettes that are located close enough to the P_C_ promoter are the only ones to be expressed, except for which that contains their own promoter (Biskri and Mazel, 2003; da Fonseca and Vicente, 2012; Stokes and Hall, 1991; Szekeres et al., 2007). The integron system represents a low-cost and modular reservoir of adaptive functions for their host bacteria. Each gene cassette is flanked by an *attC* recombination site that is recognized and recombined by the integrase. The latter catalyzes the different reactions that lead to cassette mobilization. By recombining *attI* and *attC* sites, the integrase allows the recruitment of cassettes downstream of the P_C_ promoter and express them (Figure 1A). By recombining two consecutive *attC* sites, the integrase ensures the excision of one cassette. Subsequent excision and integration of a circular cassette in the first position of the array, constitute a way for a previously silent gene to be expressed (Figure 1A).

**Figure 1:**
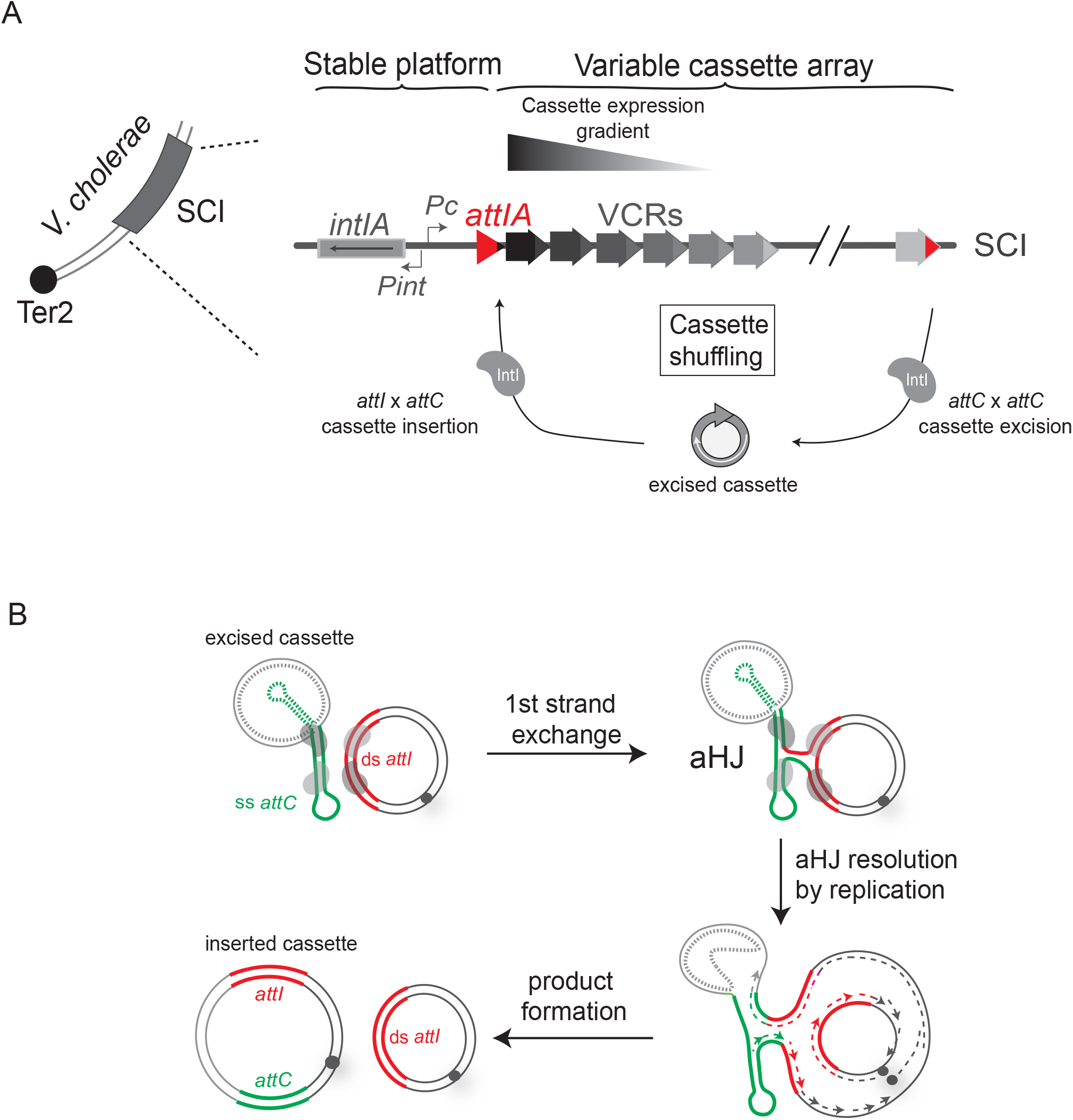
The integron. (A) *The integron system in* Vibrio cholerae. *V. cholerae* sedentary chromosomal integron is located on second chromosome close to the termination site, Ter2. The four components of integron stable platform are shown: the integrase expressing gene, *intIA*, the two promoters, P_C_ and P_int_ and the *attIA* recombination site (red triangle). The variable cassette array contains a large number of cassettes, which are represented by small arrows. Their expression level is reflected by the colour intensity of each arrow. Only the first cassettes of the array are expressed, and the subsequent ones can be seen as a low-cost cassette reservoir. Upon expression of the integrase (grey forms) cassette shuffling can occur through cassette excision (*attC* x *attC*) and integration in the first position in the array (*attIA* x *attC*). (B) *Integron cassette insertion in an* attI *site*. Recombination between the double-stranded *attI* site (bold red lines) and a single-stranded bottom *attC* site (green lines) ending a cassette is shown. Since we do not exactly know the nature of the cassettes (ss or ds), the top strand of the *attC* site is represented as a dotted line. The synaptic complex comprises both *att* sites bound by four integrase monomers (grey ovals). One strand from each *att* site is cleaved and transferred to form an atypical Holliday junction (aHJ). aHJ resolution implies a replication step. The origin of replication is represented by a grey circle and the newly synthesized leading and lagging strands by dashed lines. Both products are represented: the initial substrate resulting from the top strand replication, and the reactional product containing the inserted cassette and resulting from the bottom strand replication.

A key feature of the integron is the ability of the integrase to recombine both single-stranded DNA (*attC* site) and double-stranded DNA (*attI* site) depending on their structure and sequence respectively (Figure 1B) (Bouvier et al., 2005; Bouvier et al., 2009). Indeed, each integrase recognizes the sequence of its cognate *attI* site (Biskri et al., 2005; Collis et al., 2002). In contrast, the integrase does not recognize the sequence of *attC* sites, but rather the structure of the ss folded bottom strand (bs) (Figure 1B) (Bouvier et al., 2005; Bouvier et al., 2009; Francia et al., 1999; Johansson et al., 2004). This specific recognition of the folded bottom strand of *attC* sites allows the insertion of cassettes in the proper orientation so that they can be expressed by the P_C_ promoter (Bouvier et al., 2009; Nivina et al., 2016). The recognition of this specific ssDNA substrate imposes some constraints for recombination reactions. Indeed, during the *attI* × *attC* reaction, an atypical Holliday junction (aHJ) is formed (Figure 1B), which cannot be resolved by the classical way but can be by a host-dependent replicative pathway (Loot et al., 2012). Several other host processes are implicated in cassette recombination, for instance, by influencing the proper folding of *attC* site (Loot et al., 2010; Loot et al., 2017). Host cells also control integrase and cassette expressions (Baharoglu et al., 2012; Krin et al., 2014; Strugeon et al., 2016). The most relevant regulatory network is the induction of integrase expression, for both Class 1 MI and *V. cholerae* SCI systems, in response to environmental stress through the SOS response (Cambray et al., 2011; Guerin et al., 2009). Such regulation allows the conditional reshuffling of cassettes, at moment where the cells need to adapt to environmental changes. These examples show how integrons are intricate host-cell connected systems to maximize the potential benefit conveyed by this “adaptation on demand” device (Escudero et al., 2015).

Until then, the recombination process occurring in MIs were largely studied and the majority of assays were developed and performed in *E. coli* strains. Paradoxically, our knowledge on the SCI of the *V. cholerae* pathogenic strain remain predominantly descriptive and this, more than 20 years after its discovery and despite being the paradigm in the field. Here, for the first time, we designed experimental assays to study cassette recruitment dynamic directly in the *V. cholerae* SCI. We delivered cassettes substrates using conjugation but also natural transformation since *V. cholerae* is known to be naturally competent and to exchange DNA by this way. We obtained significant cassette insertion rate mediated by the sole endogenous integrase. We confirmed that this integrase expression is due to the SOS activation probably triggered by the single-stranded cassette delivery during conjugation and transformation processes. Interestingly, cassettes are preferentially recruited into the *attIA* primary recombination site, directly downstream of the P_C_ promoter, ensuring their expression and consecutive testing for selective advantages they can confer.

By performing *in vivo* recombination assays, we showed that RecA protein (RecA_Vch_) is critical for the cassette recruitment in the *attIA* site of the *V. cholerae* SCI. The impact of RecA_Vch_ on cassette recombination is SOS-independent and seems specific of the *attIA* × *attC* reaction mediated by the integrase of *V. cholerae*, IntIA. Indeed, the RecA_Vch_ protein did not influence neither *attC* × *attC* recombination mediated by IntIA nor the MI recombination cassettes. Moreover, unlike that of MIs, the *V. cholerae* SCI is not active in others bacterial hosts (e.g., in *E. coli,* even supplemented with the RecA_Vch_). Altogether, these results suggest that, in contrast of MIs, some specific host factors can regulate cassette recombination in SCIs. Therefore, MIs might have evolved from the SCIs by overcoming some host factors. In addition to their association with transposons and conjugative plasmids, this evolutionary trait may explain the large MI dissemination among bacteria and the antibioresistance expansion.

## MATERIAL AND METHODS

### Bacterial strains and media

*E. coli* and *V. cholerae* strains were cultivated at 37°C in Luria Bertani (LB). *V. cholerae* and *E. coli* strains, which contain a plasmid with a thermo-sensible origin of replication were grown at 30°C. Thymidine (Thy) and diaminopimelic acid (DAP) were supplemented, when necessary, to a final concentration of 0.3 mM. Glucose (Glu), L-arabinose (Ara) and isopropyl-β-D-thiogalactopyranoside (IPTG) were added in media respectively at final concentration of 10, 2 mg/ml and 0.8 mM. To induce P_tet_ promoter anhydrotetracycline (aTc) was supplemented in media to a final concentration of 1μg/ml. In the case of *E. coli* strains, antibiotics were added at the following concentrations: carbenicillin (Carb), 100 μg/ml, chloramphenicol (Cm), 25 μg/ml, kanamycin (Km), spectinomycin (Sp), 50 μg/ml. *V. cholerae* strains were cultivated with the same antibiotic concentrations except in the case of Cm and Sp, that were supplemented at a final concentration of 5 μg/ml and 100 μg/ml respectively. When *V. cholerae* strains were cultivated in presence of glucose, the later concentration of Sp was increased 2-fold (200 μg/ml).

### Plasmid and strain construction

The different plasmids and strains that were used in this study are described in Table S1 and S2. Primers used to construct the different vectors are listed in Table S3.

We performed allelic exchange to construct N16961 *ΔrecA*, *Δ*N16961 *ΔrecA ΔattIA*, *Δ*N16961 *ΔrecA ΔattIA::attI1* and N16961 *hapR+ lexA(ind−).* To this purpose, we used different variants of the pMP7 vector, respectively pB203, pK590, pK584 and p6780 (Guerin et al., 2009). We followed the same protocols as previously described (Le Roux et al., 2007; Val et al., 2012). Briefly, the suicide vector pMP7 contains a R6K origin of replication and its replication is then dependent on the presence of the Π protein in the host cell. The Π3813 cell, a *pir+* CcdB resistant *E. coli* strain (Le Roux et al., 2007), was used for cloning the different pMP7 plasmids. Once constructed, these vectors were transformed into the β3914 donor strain (Le Roux et al., 2007) in order to deliver by conjugation the pMP7 vector into the desired recipient strain. Homology regions corresponding to the genomic DNA from the recipient strain have been cloned in the different pMP7 vector to allow the integration of the plasmid by homologous recombination. The only way for pMP7 vector to replicate into recipient strains is then to integrate into the host genome after a first crossover. After conjugation, integration of the entire pMP7 vector were then selected by plating cells on Cm plates lacking DAP. Next, cells were grown in presence of L-arabinose (0.2%) in order to express the CcdB toxin. The expression of this toxin allows to kill cells in which the second crossover that leads to the excision of pMP7 backbone did not take place. This method allows us to obtain mutants of *V. cholerae* that are devoid of any antibiotic resistance marker. Note that for the deletion or replacement of *attIA* site in N16961*ΔrecA* strains, we previously transformed this cell with pAM*::recA*_*Ec*_ vector (pCY579 (Cronan, 2003). The expression of RecA_*Ec*_ protein allows allelic replacement to take place in N16961*ΔrecA* mutant. At the end of construction, strains were cultivated without Carb and IPTG and loss of the pAM*::recA*_*Ec*_ plasmid was assessed. For ectopic complementation of the *recA* mutation, we insert a copy of the *recA* gene into the *att*Tn7 site, which is present on the chromosome of *E. coli* and *V. cholerae*. We used the same strategy as the one described in (de Lemos Martins et al., 2018). The helper plasmid pMVM1 was transformed into both N16961 and MG1655 *recA* mutants. This vector has a thermo-sensitive origin of replication and carries a P_BAD_ promoter that triggers the expression of TnsABCD transposases. These transposases catalyze insertion into *att*Tn7 at high frequency. A second shuttle vector, pMP234, carries the IR sites that are recognized by the transposases and was modified for specific integration of the *recA* gene from *E. coli* or *V. cholerae*. The FRT-*aph*-FRT cassette was also added in between the IR, in order to select for transposition event. The pMP234 vector is a derivative of the pSW23T suicide vector, so its replication cannot take place into recipient cells that lack the Π protein. For integration, the pMP234 shuttle vector was delivered by conjugation in the recipient strains containing pMVM1. Transposition events were selected by plating conjugants on Km plates without DAP. These plates were incubated overnight at 42°C to get rid of the helper vector. The integration of P_LAC_-*recA*-FRT-*aph*-FRT fragment was assessed by testing UV sensitivity of the strains and by performing PCR and subsequent sequencing. After integration, the Flippase (Flp) expressing vector (pMP108, Carb^R^ (de Lemos Martins et al., 2018)) was delivered by conjugation into the *V. cholerae* strain in order to excise the Km resistance cassette. This plasmid is easily lost when culturing *V. cholerae* strains without Carb. In the case of *E. coli*, we transformed the strains with the pCP20 Flp expressing vector (Carb^R^ (Cherepanov and Wackernagel, 1995)), which has a thermo-sensitive origin replication.

### Suicide conjugation assay

This assay has been previously described (Bouvier et al., 2005) and implemented (Biskri et al., 2005) for the delivery of one specific strand of a recombination site into recipient strains that express the integrase. In this study, we used the suicide conjugative vector pSW23T (pD060) that allows the delivery of the bottom strand of the *attC*_*aadA7*_ recombination site. This vector carries a RP4 origin of transfer and an *oriV*_R6Kγ_ origin of replication. It was previously transformed into a *pir+* donor strain, β2163, which contains the RP4 machinery of transfer. This later strain needs DAP to grow in rich medium, which allows its counter-selection after that conjugation took place. The only possibility for pSW23T to replicate into the *pir−* recipient strains is then to insert into the genome through a recombination reaction catalyzed by the IntI protein. Since the pSW23T vector contains a Cm^R^ cassette, recombination events can be selected with this marker. By plating in parallel conjugants on solid media that contain or not Cm, we are able to establish the frequency of a given recombination reaction. We adapted this protocol for the use of *V. cholerae* as recipient strain, in which plasmids are more easily lost in absence of antibiotic selection than in *E. coli*. In this case, after an overnight culture, recipient cells were diluted (1:100) and grown in presence of Sp and Ara (0.2%) respectively to maintain pBAD43 vector and to allow the expression of the integrase. In the case of the *recA*_*Vch*_ and *recA*_*Ec*_ complemented strain, ITPG was also added in the media. The donor strain was grown in parallel in presence of DAP. When both donor and recipient cultures reach an OD_600nm_ of 0.6, 1ml of each culture was harvested by centrifuging 6 min at 6000 rpm. The obtained pellet was re-suspended in a droplet of LB and spread on a 0.45 μm filter. This filter was placed on MH DAP, Ara plates and incubated at 37°C during 3h. After incubation, the filter was re-suspended in 5ml of LB and this suspension was used to spread appropriate dilutions on MH, Cm, Sp, Glu and MH, Sp, Glu plates. After 2 days of incubation at 37°C, the recombination frequency was calculated as the ratio of Cm^R^ clones over the total number of recipient colonies that grew on MH, Sp, Glu plates. Note that in the case where we did not detect recombination event for one replicate, we calculate the recombination rates as the ratio of the mean of recombinant clones over the mean of total recipient clones obtained for the different replicates. In this case no error bars are represented on our graphics.

### Integrase mediated recombination after natural transformation

In this study, we used the same pSW23T vector (pD060) that used in the suicide conjugation assay. *V. cholerae* natural transformation was previously described (Marvig and Blokesch, 2010). Here, we adapted this protocol to assess integrase-mediated recombination frequency after natural transformation in naturally competent strains (*hapR+*) of *V. cholerae*. An overnight culture was used to inoculate 1:100 of bacteria in 5ml of LB medium supplemented with Sp (50 μg/ml). The culture was grown at 30°C until they reached an OD_600nm_ of 0.5. One milliliter of cells was then centrifuged (5000 rpm, 10 minutes) and resuspended in 1 ml of M9 minimal medium supplemented with MgSO4 (32 mM) and CaCl_2_ (5 mM) and Sp (100 μg/ml). Tubes containing 50 to 80 mg of chitin (C9213; Sigma) were inoculated with 0.5 ml of washed cells and 0.5 ml of fresh M9 medium supplemented with MgSO4, CaCl_2_ and Sp, vortexed, and grown 48 hours at 30°C with shaking. Cultures were then washed (5000 rpm, 10 minutes) and resuspended in an equal volume of fresh M9 medium supplemented with MgSO4, CaCl_2_, Sp and anhydrotetracycline (aTc) to induce integrase expression. The cultures were incubated again for 30 minutes at 30°C with shaking. Then, 2 μg of plasmid DNA (the pD060 vector) were added to the cultures and incubated for 36 h at 30°C with shaking. After incubation, bacteria were detached from chitin by vortexing vigorously during 30 seconds. The obtained bacteria suspension was used to spread appropriate dilutions on MH, Cm, Sp and MH, Sp plates. After 2 days of incubation at 37°C, the recombination frequency was calculated as the ratio of Cm^R^ clones over the total number of recipient colonies that grew on MH, Sp plates in the same manner as for suicide conjugation assay.

### Analysis of cassette insertion point localization

For each experiment, at least eight recombinant clones were isolated on appropriate plates and analyzed by PCR. For this, we performed different PCR reactions. In order to determine precisely if the pSW23T vector has been inserted into the *attIA* site of the SCI we used 5778 and SWend primers. These primers hybridize respectively in a sequence upstream of *attIA* in *V. cholerae* chromosome 2 or downstream of *attC*_*aadA7*_ in the pSW23T vector. For *E. coli* and *V. cholerae* strains transformed with the pSU38Δ or on pBAD43 vectors that harbor recombination sites, SWbeg and MFD or 1704 and SWend primers respectively were used to amplify one junction of the co-integrate. Finally, to detect insertion of pSW23T in the genome of *V. cholerae*, either at secondary sites or into the VCR sites of the SCI, we performed random PCR amplification. For these, we performed a first random PCR reaction using the 1863 degenerated and the 2405 primers. The 2405 primer hybridizes upstream of the *attC* sites on pSW23T plasmids. Due to the presence of degenerate nucleotides in the 1863 primer, low hybridization temperatures were used, first, 30°C during 5 cycles and after, 40° during 30 cycles. The obtained amplified DNA fragments were subjected to a second PCR reaction in order to enrich for PCR products corresponding to cassette insertion. For this purpose, we used 1865 and 1388 primers. These primers hybridize respectively to the fixed part of the degenerated 1863 primer and upstream (but closer than 2405) of the *attC* sites on pSW23T plasmids. Recombination points were precisely determined by sequencing PCR products using 1366. The 1366 primer hybridizes upstream (but closer than 1388) of the *attC* sites on pSW23T plasmids. In all cases, some PCR reactions were purified and sequenced to confirm the insertion point.

### Recombination assay with unidirectional replicative substrate

This assay was previously described (Loot et al., 2010) and allow the determination of the recombination rates of a given recombination reaction when *attC* sites are carried by a replicative plasmid. The vectors that we used (p7523 or p7546, Cm^R^) replicate unidirectionally and the *attC*_*aadA7*_ recombination sites they carry have been cloned, so that the recombinogenic bottom strand is located either on leading or lagging strand template (“lead” or “lag” orientation). *V. cholerae* strains were transformed with these unidirectional-replicative vectors and the IntIA expressing vector (p995, Carb^R^) or with the empty version of the plasmid (p979, Carb^R^). As p7523 and p7546 have both a thermo-sensitive origin of replication, the cells were cultivated overnight at 30°C. The next day, cultures were diluted (1:100) and incubated during 5h at 30°C in presence of Cm and Carb to maintain both pTSC29 and pBAD18 plasmid. In order that recombination takes place, L-arabinose was also added in the media at a concentration of 0.2% to allow the expression of the integrase. After 5h of incubation, appropriate dilutions of the cultures were plated in parallel on MH Cm, Carb, Glu and MH Carb, Glu plates and incubated two days at 42°C. Since pTSC29 vector cannot replicate at 42°C, the selection on Cm plates at this temperature allows recovering only the clones in which *attC*_*aadA7*_ site have been recombined by IntIA. Recombination rates were calculated as for suicide conjugation assays, by considering the ratio of Cm^R^ clones over total number of recipient clones that grew on MH Carb, Glu plates. In order to determine if cassette insertion occurs into the *attIA* site of the SCI of *V. cholerae*, we performed PCR reactions on at least eight random isolated clones using 5778 and 573 or 5778 and MFD primers (respectively used for “lead” and “lag” orientation of the bs of *attC*_*aadA7*_ site on pTSC vector).

## RESULTS

### *Horizontally transferred cassettes are efficiently inserted into the* attIA *site of the* Vibrio cholerae *SCI*

Large SCIs such as the *V. cholerae* SCI constitute huge reservoirs of functions for their host bacteria. In such large integrons, several recombination sites can constitute potential targets for cassette insertion (*attIA* primary recombination site and/or the numerous VCR sites). In a previous study, recombination assays that were performed in *V. cholerae* aimed at evaluating cassette insertion in recombination sites carried on plasmids (Biskri et al., 2005). Here, we attempt to visualize cassette insertion into the SCI of *V. cholerae* using our classical conjugation assay (Bouvier et al., 2005) and developing a natural transformation assay. Interestingly, both assays reproduce the natural conditions in which the acquisition of cassettes could occur through horizontal gene transfer in *V. cholerae* (Figure 2A). The plasmids containing the donor *attC* site (pSW23T*::attC*_*aadA7*_) are delivered on a single-stranded form in a recipient *V. cholerae* strain containing a vector expressing the IntIA integrase. Once delivered, *attC*-containing plasmids cannot replicate and can therefore be assimilated to a non-replicative integron cassette. The unique way for these synthetic cassettes to be maintained is to be inserted in the *V. cholerae* host genome by a recombination between the cassette *attC* sites and the SCI *att* sites.

**Figure 2:**
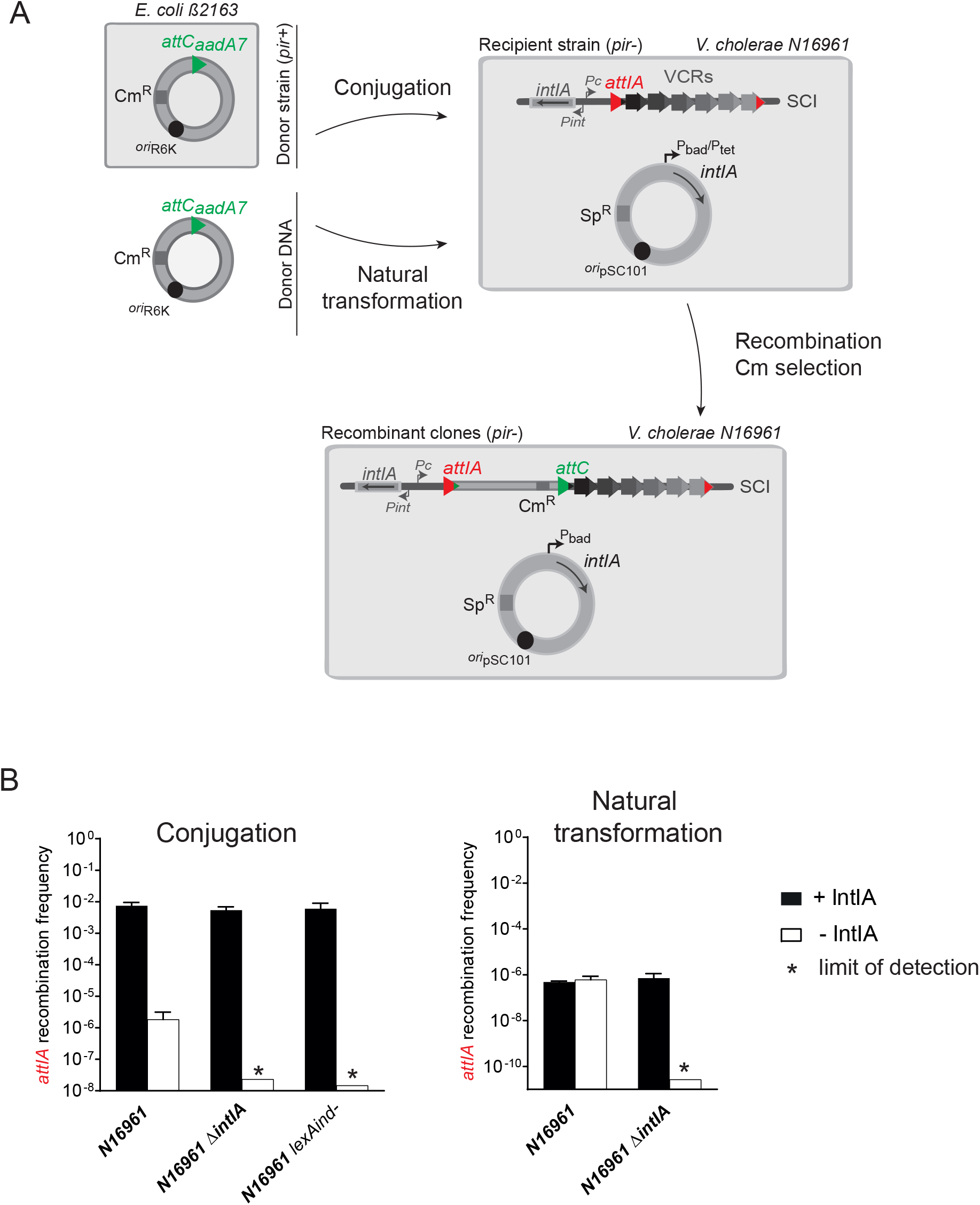
Cassette recruitment in *V. cholerae* SCI during horizontal gene transfer. (A) Experimental setup of the cassette insertion assay The pSW23T::*attC*_*aadA7*_ suicide vector is delivered to N16961 *V. cholerae* recipient strains containing an integrase expressing vector or the sole endogenous integrase, and the SCI. The delivering occurs by two horizontal gene transfer processes: conjugation from the β2163 donor or natural transformation. As the pSW23T cannot replicate in *V. cholerae* recipient strains, recombinant clones can be selected on appropriate Cm containing plates to evaluate the recombination frequency (see also Results and Material and methods). The *attC*_*aadA7*_ site carried by the suicide vector is represented by a green triangle and the *attIA* site on the *V. cholerae* SCI by a red triangle. (B) Frequency of insertion of the pSW23T*::attC*_*aadA7*_ suicide vector into the *attIA* site. The recombination frequencies were calculated in N16961 *V. cholerae* wt, *Δ*intIA *Δ*and *lexAind-* strains. Results correspond to recombination frequencies that were normalized after analysis of PCR reactions (Material and Methods). +IntIA: recipient strains transformed with the pBAD43 integrase expressing vector; −IntIA: control strains transformed with the empty pBAD43 vector. * correspond to the limits of detection. Values represent the mean of at least three independent experiments and error bars correspond to average deviations from the mean.

When we carried out this test by delivering cassette by conjugation assay, we detected a significant level of insertion (7.5 × 10^−3^) of the pSW23T*::attC*_*aadA7*_ vector using the N16961 *wt V. cholerae* strain (Figure 2B, + IntIA). We performed PCR reactions on some recombined clones and demonstrated that, in all cases, the cassette insertion occurs at *attIA* sites from *V. cholerae* SCI platform (Figure S1A). By sequencing some PCR products, we confirmed that insertion point was correctly localized in the 5’-AAC-3’ triplet.

We took advantage from *V. cholerae* natural competent state in the presence of chitin (Meibom et al., 2005) to investigate for cassette recombination in the context of another HGT mode. We adapted natural transformation protocol to evaluate cassette insertion frequency in the *V. cholerae* SCI (Figure 2A). We obtained a recombination frequency of 4.8 × 10^−7^ when overexpressing the integrase (Figure 2B, +IntIA). Importantly, these recombination frequencies, which are much lower than those obtained during conjugation in the same conditions (integrase overexpression), are the consequences of a limited proper frequency of natural transformation (Marvig and Blokesch, 2010). We also performed PCR reactions on some recombined products and demonstrated that cassette insertion occurs, for all tested clones, at *attIA* sites from *V. cholerae* SCI platform (Figure S1B). These results show that, during conjugation or natural transformation, integron cassettes are efficiently released in the *V. cholerae* host cell and inserted in the SCI. Interestingly, the large majority of insertion events occurs in the integron platform *attIA* in spite of the presence of 180 *attC* sites meaning that all newly inserted cassettes are expressed and therefore tested for their selective advantage.

### *The endogenous integrase efficiently inserts cassettes into the* attIA *site of the* Vibrio cholerae *SCI*

We also performed the both assays in the presence of the sole endogenous SCI IntIA integrase. Interestingly, in this case, we also detected a significant level of insertion of the pSW23T*::attC*_*aadA7*_ vector using the N16961 *wt V. cholerae* strain (1.8 × 10^−6^ and 6.1 × 10^−7^ respectively for conjugation and transformation assays, Figure 2B, −IntIA). Here again, we performed PCR reactions on some recombined products and demonstrated that cassette insertion occurs, for all tested clones, at *attIA* sites from *V. cholerae* SCI platform. This recombination activity is due to the expression of the endogenous *intIA* integrase gene since no recombination event was detected in the strain devoid of endogenous integrase (N16961 *ΔintIA*) for both assays (Figure 2B, −IntIA). We also demonstrated that the expression of endogenous integrase is dependant of the SOS system since no recombination event was detected below the detection limit of 1.5 × 10^−8^ in the N16961 *lexAind-* strain in which the SOS response is not inducible. Indeed, in this strain, the SOS regulon genes are constitutively repressed because of the presence of the uncleavable LexA_A91D_ version of the LexA repressor (Guerin et al., 2009). As supplementary control, we also overexpressed the IntIA integrase in these both mutant strains and as expected we obtained a very high recombination frequency (respectively 5.4 × 10^−3^ and 6.0 × 10^−3^ in the *ΔintIA* and *lexAind-* strains, Figure 2B, + IntIA). Altogether, these results show that, during conjugation or natural transformation, integron cassettes are efficiently released in the *V. cholerae* host cell and inserted in the SCI even in the presence of the sole endogenous integrase. The level of endogenous integrase expression, triggered by SOS response induction initiated by the single-stranded cassette entry during conjugation and natural transformation (Baharoglu et al., 2010) seems sufficient to insert cassettes at significant level in the *V. cholerae* SCI. Note that, when performing both conjugation and natural transformation, we detected some *attIA* insertion events associated with shuffling of cassettes in first position (1/160 and 12/152 respectively, Figure S1).

### *RecA*_*Vch*_ *influences cassette insertion into the* attIA *site of the* Vibrio cholerae *SCI*

Here, we precisely investigated the cassette recombination mechanism used by the SCI of *V. cholerae*. To define the network of intervening host factors, we tested the effect of the *Vibrio cholerae* RecA protein (RecA_Vch_). The RecA protein is functionally conserved among bacterial species (Goldberg and Mekalanos, 1986) and in eukaryotic organisms (Shinohara and Ogawa, 1999). RecA is a critical enzyme for homologous recombination process, during which it binds ssDNA catalyzing the pairing with complementary regions of dsDNA and strand exchange reactions (Cox, 2007a; Kowalczykowski, 2000; Lusetti and Cox, 2002). Because of the capacity of the RecA protein to bind ssDNA, we tested its impact on SCI cassette recombination catalyzed by IntIA in *V. cholerae*.

Among the several conditions previously developed, we chose to use the optimal one, i.e., the suicide conjugation assay with overexpression of integrase (Figure 3A). Here again, we obtained a very high recombination rate (2.0 × 10^−2^) when using the *wt* parental strain as recipient strain. We observed a decrease of more than two orders of magnitude in the recombination rates (7.7 × 10^−5^) in the corresponding N16961 *ΔrecA* mutant strain (Figure 3A). When performing PCR analysis for each reaction, we confirmed that insertions occur in the *attIA* site (222/222 and 108/111, respectively for the *wt* and *ΔrecA* strains). These results mean that the RecA_*Vch*_ protein favors insertion of cassettes in the *attIA* site of *V. cholerae* SCI. As expected, no recombination event was detected in the N16961 *ΔrecA* control strain that carries the empty pBAD43 vector (compare to the N16961 *wt* strain, Figure 3A), since the SOS response cannot be activated and induce the endogenous integrase expression in these cells. To confirm that the decrease in recombination rates that we observed is specifically due to the deletion of the *recA*_*Vch*_ gene and not to polar effects, we constructed a strain in which the *recA*_*Vch*_ deletion was ectopically complemented. For this, a copy of the *recA*_*Vch*_ gene was inserted into the *att*Tn7 site located in the chromosome 1 of *V. cholerae* (de Lemos Martins et al., 2018). The *recA*_*Vch*_ ectopic complementation allows recovering the same recombination rate as in the N16961 *wt* strain (Figure 3A), meaning that the effect on *attIA* × *attC*_*aadA7*_ recombination reaction, which we observed, is specific to the absence of the RecA_*Vch*_ protein. Furthermore, we tested the ability of the RecA protein of *E. coli* (RecA_*Ec*_) to complement the *recA*_*Vch*_ deletion. The RecA protein of *E. coli* and *V. cholerae* show variations in the amino acid sequence of their respective C-terminal part (Figure S2). This C-terminal region is implicated in the modulation of RecA_*Ec*_ activity notably by interacting with regulator proteins (Cox, 2007b; Lusetti et al., 2004). We then tested if such variations could lead to host specific regulation that will impact cassette recombination. For this, we inserted a copy of the *recA* gene of *E. coli* into the *att*Tn*7* locus in *V. cholerae*. We therefore demonstrated that the complementation of the N16961 *ΔrecA* mutant with RecA_*Ec*_ was as efficient as with the native RecA_*Vch*_ of *V. cholerae* (Figure 3A). This is consistent with the high percentage of identity (almost 80 %, Figure S2) shared by both RecA proteins. Moreover, we observed that the expression of RecA_*Ec*_ also triggers SOS induction leading to endogenous integrase expression in the control strain harboring the empty pBAD43 plasmid (Figure 3A). Both RecA proteins seem then to share enough identity so that the RecA_*Ec*_ can replace RecA_*Vch*_ for several functions including its apparent role in cassette recombination in *V. cholerae*.

**Figure 3:**
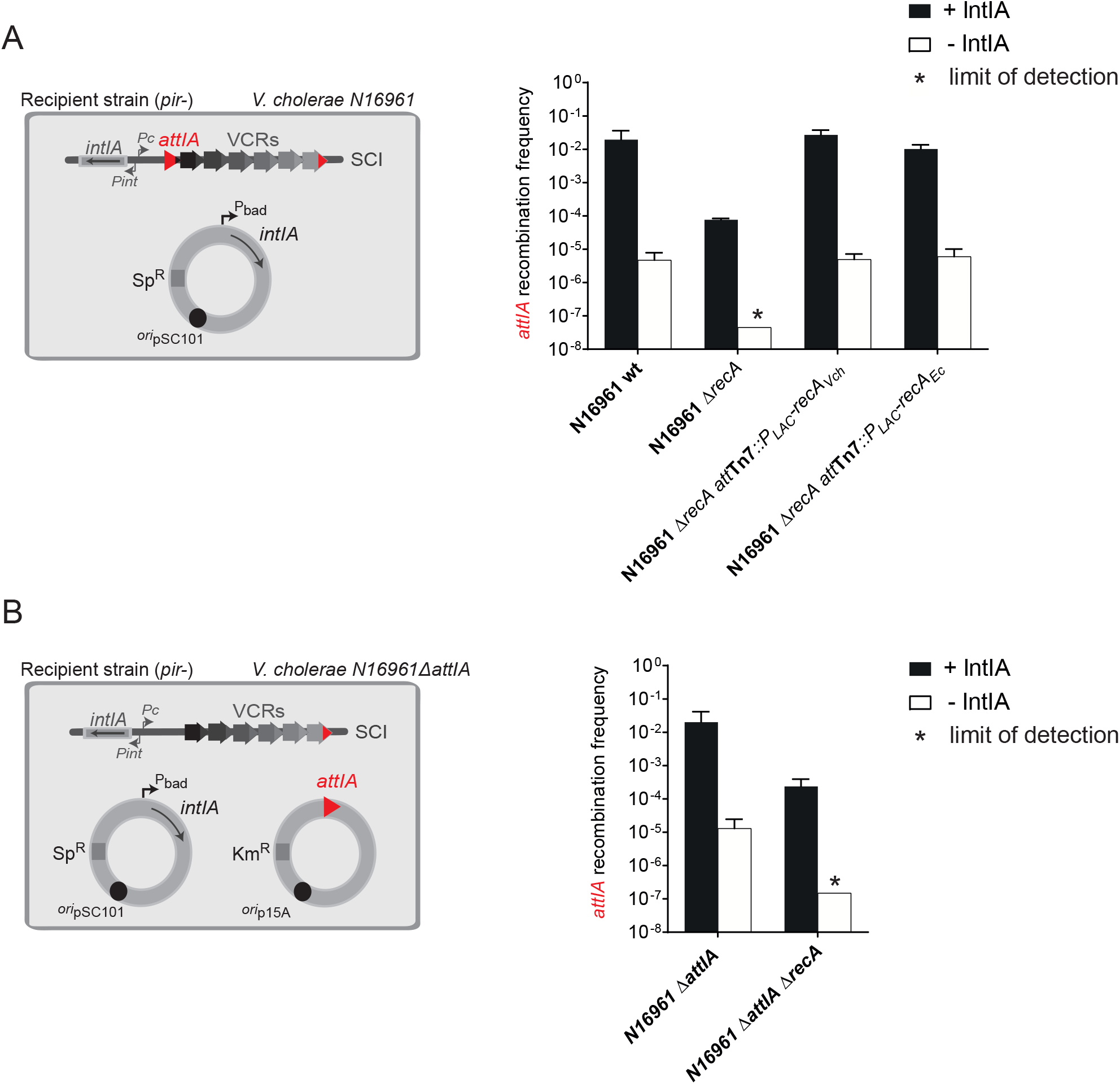
Effect of the RecA protein on *attIA* × *attC* recombination in *V. cholerae* SCI. (A) Experimental setup and frequency of insertion of the pSW23T*::attC*_*aadA7*_ suicide vector into the chromosomic *attIA* site. N16961 recipient strains transformed with the pBAD43 IntIA expressing vector were used (left panel). The recombination rates were calculated in N16961 *V. cholerae* wt and in the corresponding *recA* mutant (*ΔrecA*) and ectopic complemented (*Δ*recA-att_tn7_::*Δ*P_LAC_-*recA*_*Vch*_ and *Δ*recA-att_tn7_::*Δ*P_LAC_-*recA_Ec_*) strains (right panel). (B) Experimental setup and frequency of insertion of the pSW23T*::attC*_*aadA7*_ suicide vector into the *attIA* site located on plasmid. N16961 recipient strains transformed with both pBAD43 IntIA expressing vector and pSU38Δ*::attIA* vector were used (left panel). The recombination rates were calculated in N16961 *V. cholerae* wt and in the corresponding *recA* mutant strains (*ΔrecAΔ*, right panel). For both (A) and (B), results correspond to recombination frequencies that were normalized after analysis of PCR reactions (Material and Methods). +IntIA: recipient strains transformed with the pBAD43 integrase expressing vector; −IntIA: control strains transformed with the empty pBAD43 vector. * correspond to the limits of detection. Values represent the mean of at least three independent experiments and error bars correspond to average deviations from the mean.

Since RecA is involved in the induction of the SOS response, its effect on cassette recombination could be indirect and involve one or more proteins of the SOS regulon. Yet, we previously demonstrated that into the *lexAind-* mutant, in which the SOS response is non-inducible, we did not observe any effect on recombination efficiency compare to the *wt* strain (Figure 2B). The use of this strain allows us to uncouple the effect on SOS induction from other properties of the RecA protein and to demonstrate that the RecA_*Vch*_ effect is not due to the SOS regulon proteins. We also determined the role of the RecA_*Vch*_ protein during cassette recombination in *attIA* when this site is moved on a plasmid (Figure 3A). For this, we constructed a *V. cholerae* strain deleted for the resident *attIA* site (*V. cholerae* N16961 *ΔattIA*) and transformed with an *attIA*-containing plasmid. Once again, we obtained a decrease of two orders of magnitude (from 2.0 × 10^−2^ to 2.4 × 10^−4^, Figure 3B).

Together these results indicate that the RecA protein of *V. cholerae* favors the *attIA* × *attC* recombination mediated by IntIA, in a SOS-independent manner and whatever the *attIA* localization.

### *RecA*_Vch_ *does not influence* attC × attC *recombination in* Vibrio cholerae

During both *attC* × *attC* and *attI* × *attC* reactions, the proper folding of *attC* site is required for binding of integrase monomer. If the RecA protein acts on cassette recombination through its ssDNA binding capacity affecting *attC* site folding, we expect that it will impact both *attI* × *attC* and *attC* × *attC* reactions that are catalyzed by IntIA. To test this, we used different assays that allow to assess cassette insertion frequencies directly into the VCR sites of the SCI or in *attC* sites carried on plasmids (Figure 4). To observe insertion events into the VCR sites of the SCI, we used the *V. cholerae* mutant strain devoid of the resident *attIA* site (*V. cholerae* N16961 *ΔattIA*, see just above). Here, we did not observe any effect of the RecA protein in the recombination rates obtained with *ΔattIA* and *ΔattIA ΔrecA*_*Vch*_ mutant strains. To confirm the cassette insertion in VCRs, we performed random PCR reactions (Material and methods). The recombination rates into VCR sites is similar between both mutant strains (Figure 4A) meaning that the RecA_*Vch*_ protein does not affect the *attC* × *attC* recombination.

**Figure 4:**
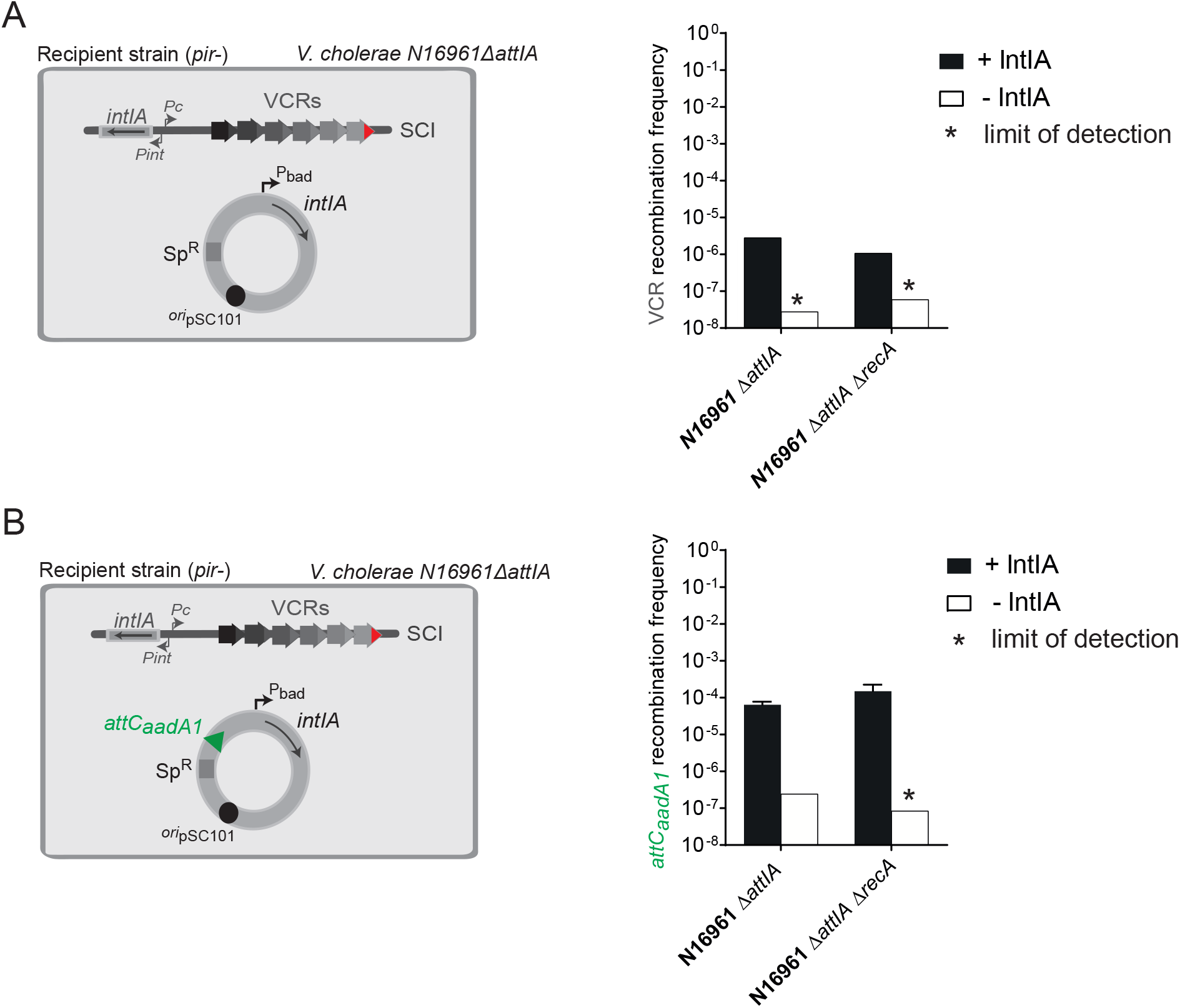
Effect of the RecA protein on *attC* × *attC* recombination in *V. cholerae*. (A) Experimental setup and frequency of insertion of the pSW23T*::attC*_*aadA7*_ suicide vector into VCR sites of the SCI. N16961 recipient strains deleted for the *attIA* site (*Δ*attIA *Δ*strains) and transformed with the pBAD43 IntIA expressing vector were used (left panel). The recombination rates were calculated in N16961 *V. cholerae *Δ*attIA* and in the corresponding *recA* mutant strain (*ΔattIA ΔrecA*, right panel). (B) Experimental setup and frequency of insertion of the pSW23T::*attC*_*aadA7*_ suicide vector into the *attC*_*aadA1*_ site located on the pBAD43 plasmid. N16961 recipient strains deleted for the *attIA* site (*ΔattIA* strains) and transformed with the IntIA expressing and *attC*_*aadA1*_ containing pBAD43 vector were used (left panel). The recombination rates were calculated in N16961 *V. cholerae ΔattIA* and the corresponding *recA* mutant strain (*ΔattIA ΔrecA*, right panel). For both (A) and (B), results correspond to recombination frequencies that were normalized after analysis of PCR reactions and sequencing (Material and Methods). +IntIA: recipient strains transformed with the pBAD43 integrase expressing vector; −IntIA: control strains transformed with the empty pBAD43 vector. * correspond to the limits of detection. Values represent the mean of at least three independent experiments and error bars correspond to average deviations from the mean.

To confirm this absence of RecA_*Vch*_ protein effect on *attC* × *attC* recombination, we perform a second conjugation assay in *V. cholerae* N16961 *ΔattIA* strains transformed with a plasmid carrying the MI *attC*_*aadA1*_ site (from the *aadA1* gene cassette). We obtained a significant rate of recombination for both *ΔattIA* and *ΔattIA ΔrecA*_*Vch*_ strains and we did not observe any impact of the RecA_*Vch*_ protein on *attC*_*aadA1*_ × *attC*_*aadA7*_ reaction catalyzed by IntIA in *V. cholerae* (Figure 4B). Together, these results, with results presented in the last paragraph, indicate that the RecA_*Vch*_ protein seems to favor only *attIA* × *attC* reaction and not to influence insertion reactions into *attC* sites, whether they are present on the chromosome (VCR) or on a plasmid (MI *attC* sites).

### *RecA*_*Vch*_ *does not influence* attI1× attC *mediated by IntI1 in* Vibrio cholerae

The previously demonstrated influence of the RecA_*Vch*_ protein on cassette recombination was striking since it was already demonstrated that the RecA_*Ec*_ protein does not influence the recombination reactions catalyzed by IntI1 in *E. coli* (Loot et al., 2014). To determine if the role of the RecA_*Vch*_ protein is specifically linked to *V. cholerae* SCI system, we used two different recombination tests to establish, in *V. cholerae*, the activity of the integrase IntI1. First, we performed the suicide conjugation assays in a *V. cholerae* strain where the *attIA* site from the SCI platform was replaced by the *attI1* site (the cognate site of the IntI1 integrase). Secondly, we performed this assay in *V. cholerae* strains deleted for the *attIA* site (*V. cholerae* N16961 *ΔattIA*) and transformed with an *attI1*-carrying plasmid. We observed, using these two assays, that the frequencies of *attI1* × *attC* reaction catalyzed by IntI1 were the same for both *ΔattIA* and *ΔattIA ΔrecA* recipient strains (Figure 5A and 5B). Therefore, we demonstrated that, in *V. cholerae*, the *attI1* × *attC* reaction catalyzed by IntI1 is not influenced by RecA_*Vch*_. This suggests that the RecA_*Vch*_ influence of the *attI* × *attC* reaction is specific to IntIA.

**Figure 5:**
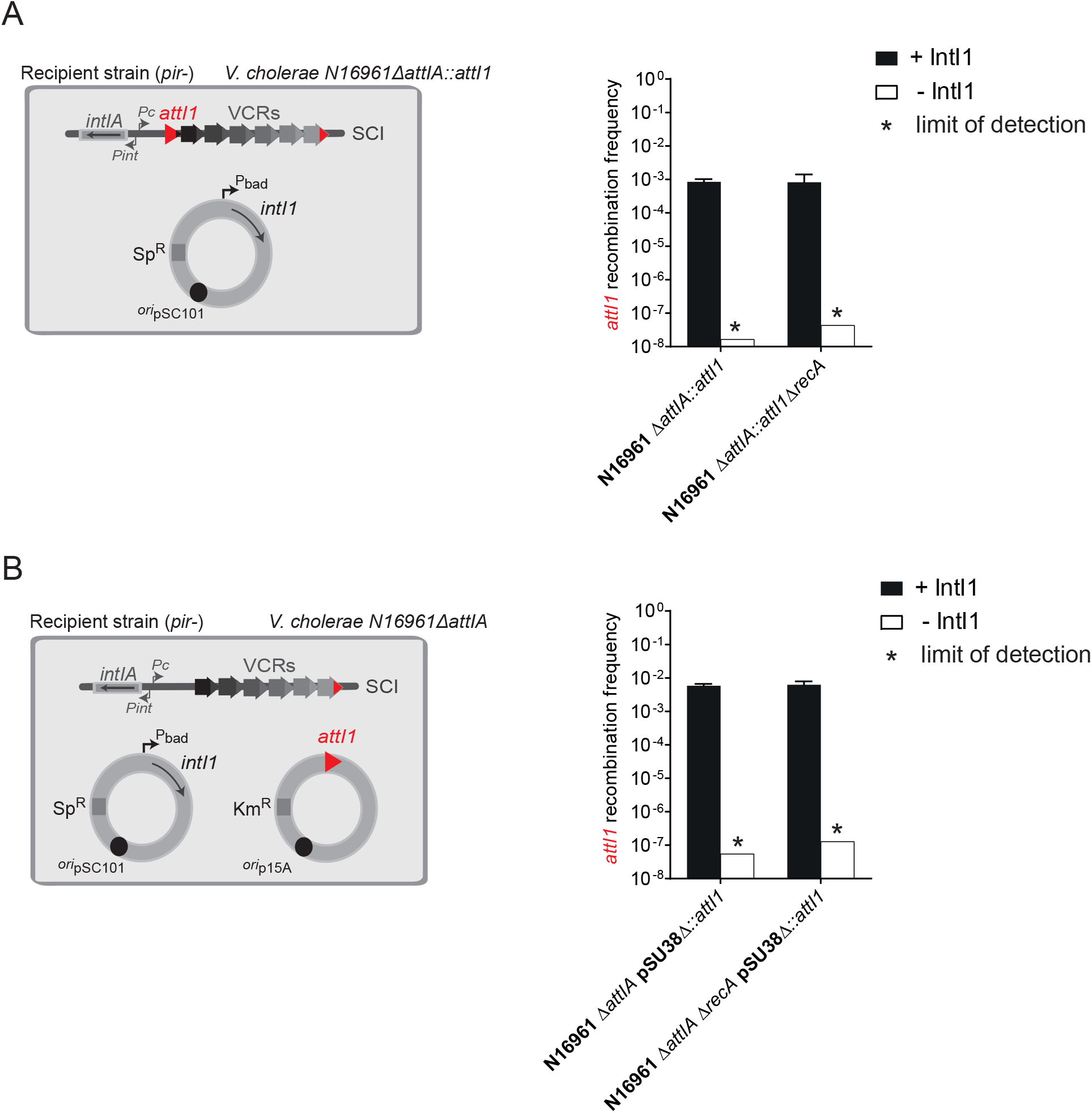
Effect of the RecA protein on *attI1* × *attC* recombination mediated by IntI1 in *V. cholerae*. (A) Experimental setup and frequency of insertion of the pSW23T*::attC*_*aadA7*_ suicide vector into the *attI1* site located in the SCI platform. N16961 recipient strains with an *attI1* site in place of the *attIA* site (*ΔattIA::attI1* strains) were transformed with the pBAD43 IntI1 expressing vector (left panel). The recombination rates were calculated in N16961 *V. cholerae ΔattIA::attI1* and the corresponding *recA* mutant strain (*ΔattIA::attI1 ΔrecA*, right panel). (B) Experimental setup and frequency of insertion of the pSW23T*::attC*_*aadA7*_ suicide vector into the *attI1* site located on a plasmid. N16961 recipient strain deleted for the *attIA* site (*ΔattIA::attI1* strain) and transformed with both pBAD43 IntI1 expressing vector and pSU38Δ*::attI1* vector were used (left panel). The recombination rates were calculated in *V. cholerae *Δ*attIA::attI1* and the corresponding *recA* mutant strain (*ΔattIA::attI1 ΔrecA*, right panel). For both (A) and (B), results correspond to recombination frequencies that were normalized after analysis of PCR reactions (Material and Methods). +IntIA: recipient strains transformed with the pBAD43 integrase expressing vector; −IntIA: control strains transformed with the empty pBAD43 vector. * correspond to the limits of detection. Values represent the mean of at least three independent experiments and error bars correspond to average deviations from the mean.

### *Effect of RecA on cassette recombination in* Escherichia coli

As already introduced, the impact of the RecA protein on cassette recombination was already tested in *E. coli* (Loot et al., 2014). In our previous study, a setup based on the reconstitution of a functional *dapA* gene after cassette excision through an intramolecular reaction was used. In this case, only the efficiency of reactions catalyzed by the integrase of Class 1 MI was studied (Loot et al., 2014). Here, we assessed the efficiency of either *attI* × *attC* or *attC* × *attC* intermolecular reactions and we tested the effect of *recA* deletion on reactions catalyzed by both IntI1 and IntIA integrases. We also used another *attC* site, VCR_VCA0441_ (Loot et al., 2014). We still observed the RecA-independence of IntI1 for both *attI1* × *attC*_*aadA7*_ and VCR × *attC*_*aadA7*_ reactions (Figure 6A and 6B, left panels). Indeed, respective recombination frequencies were identical in MG1655 *wt* and *ΔrecA* recipient strains. For the reaction catalyzed by IntIA, we observed that, in *E. coli*, the VCR × *attC*_*aadA7*_ reaction was not affected by the deletion of the *recA* gene (Figure 6B, right panel). In accordance with the study of Biskri and coll. (Biskri et al., 2005), we found that IntIA_*Vch*_ was not able to efficiently catalyze the *attIA* × *attC*_*aadA7*_ reaction in *E. coli* in the *wt* strain (Figure 6A, right panel). As proposed earlier, if IntIA recombines much less efficiently in *E. coli* this may reflect the absence, or a too large divergence, of at least one host factor required for this reaction. Thus, in order to determine if RecA could correspond to one of these divergent factors, we tested if RecA_*Vch*_ could rescue the lack of *attIA* × *attC* recombination in *E. coli*. We inserted, as performed previously in *V. cholerae*, the *recA*_*Vch*_ gene into the unique *att*Tn*7* site of *E. coli*. However, we found that the expression of RecA_*Vch*_ do not restore the recombination capacity of IntIA for the *attIA* × *attC*_*aadA7*_ reaction in *E. coli*. Thus, it shows that the RecA protein is not the factor whose divergence hamper the *attIA* × *attC*_*aadA7*_ reaction catalyzed by IntIA to take place in *E. coli*.

**Figure 6:**
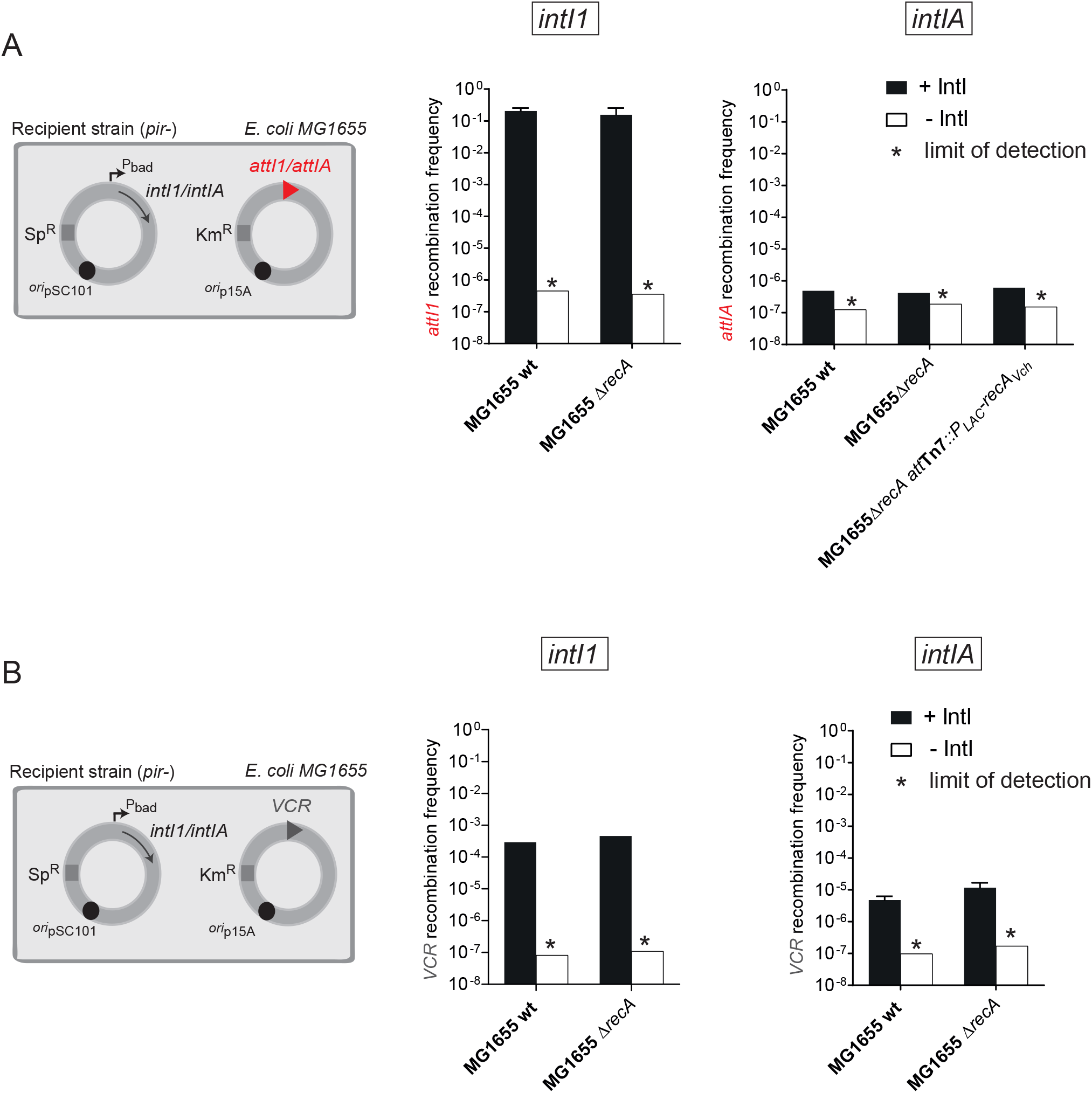
Effect of the RecA protein on recombination reactions catalysed by IntI1 and IntIA in *E. coli.* (A) Experimental setup and frequency of insertion of the pSW23T*::attC*_*aadA7*_ suicide vector into the *att* sites (*attI1* or *attIA*) located on a plasmid. MG1655 recipient strains transformed with both pBAD43 IntI expressing vector (IntI1 or IntIA) and the pSU38Δ*::attI* vector (*attI1* or *attIA*) were used (left panel). The recombination rates were calculated in MG1655 and in the corresponding *recA* mutant (*ΔrecA*) and ectopic complemented (*ΔrecA-att*_*tn7*_::P_LAC_-*recA*_*Vch*_) strains (right panels). (B) Experimental setup and frequency of insertion of the pSW23T*::attC*_*aadA7*_ suicide vector into the VCRVCA0441 site located on a plasmid. MG1655 recipient strains transformed with both pBAD43 IntI expressing vector (IntI1 or IntIA) and the pSU38Δ::VCR_VCA0441_ vector were used (left panel). The recombination rates were calculated in MG1655 and in the corresponding *recA* mutant strain (*ΔrecA*, right panels). For both (A) and (B), results correspond to recombination frequencies that were normalized after analysis of PCR reactions (Material and Methods). +IntI: recipient strains transformed with the pBAD43 integrase expressing vector; -IntI: control strains transformed with the empty pBAD43 vector. * correspond to the limits of detection. Values represent the mean of at least three independent experiments and error bars correspond to average deviations from the mean.

### *RecA*_*Vch*_ *influences* attIA *cassette insertion however the* attC *site is delivered*

In all previously presented assays, we used horizontal gene transfer mechanisms as a mean for delivering the single-stranded recombinogenic bottom strand of the *attC* site on non-replicative substrate mimicking integron cassettes. To determine if RecA still favors cassette insertion in the case where *attC* sites are folded from double-stranded molecules, we performed a recombination assay that relies on the use of replicative vectors (pTSC29 derivatives). These vectors replicate unidirectionally and the *attC*_*aadA7*_ site that they carry was cloned in two orientations, so that the bottom strand of the *attC* site is either present on lagging or leading strand template (Figure 7A). When the bs of *attC* site is located on the lagging strand template, in which large region of ssDNA are available during replication (i.e., between Okazaki fragments), bs *attC* site folding from ssDNA is favored (Figure 7B). When bs of the *attC* site is located on the leading strand template, bs *attC* site folding can only occur from dsDNA, by cruciform structures extrusion (Loot et al., 2010) (Figure 7B). Since the pTSC29 vectors have a thermo-sensitive origin of replication, the selection of recombinant Cm^R^ clones at 42°C, allows us to evaluate the efficiency of recombination events. When we performed this replicative assay in the *V. cholerae* N16961 *wt* strain, we observed a decrease of about one order of magnitude when the *attC*_*aadA7*_ bs is present on the leading strand template (Figure 7C). Such slight differences in the recombination frequency between the two orientations of the *attC*_*aadA7*_ site were previously observed, when the same recombination assays were carried out in *E. coli* with the integrase IntI1 (Loot et al., 2010). Concerning the N16961 *ΔrecA* mutant, we still observed a large decrease in recombination rates for both *attC*_*aadA7*_ site orientations (Figure 7C). These results demonstrate that RecA_*Vch*_ is involved in the mechanism of recombination during *attIA* × *attC* reaction whatever the manner that the *attC* site is delivered (i.e., from ssDNA or dsDNA).

**Figure 7:**
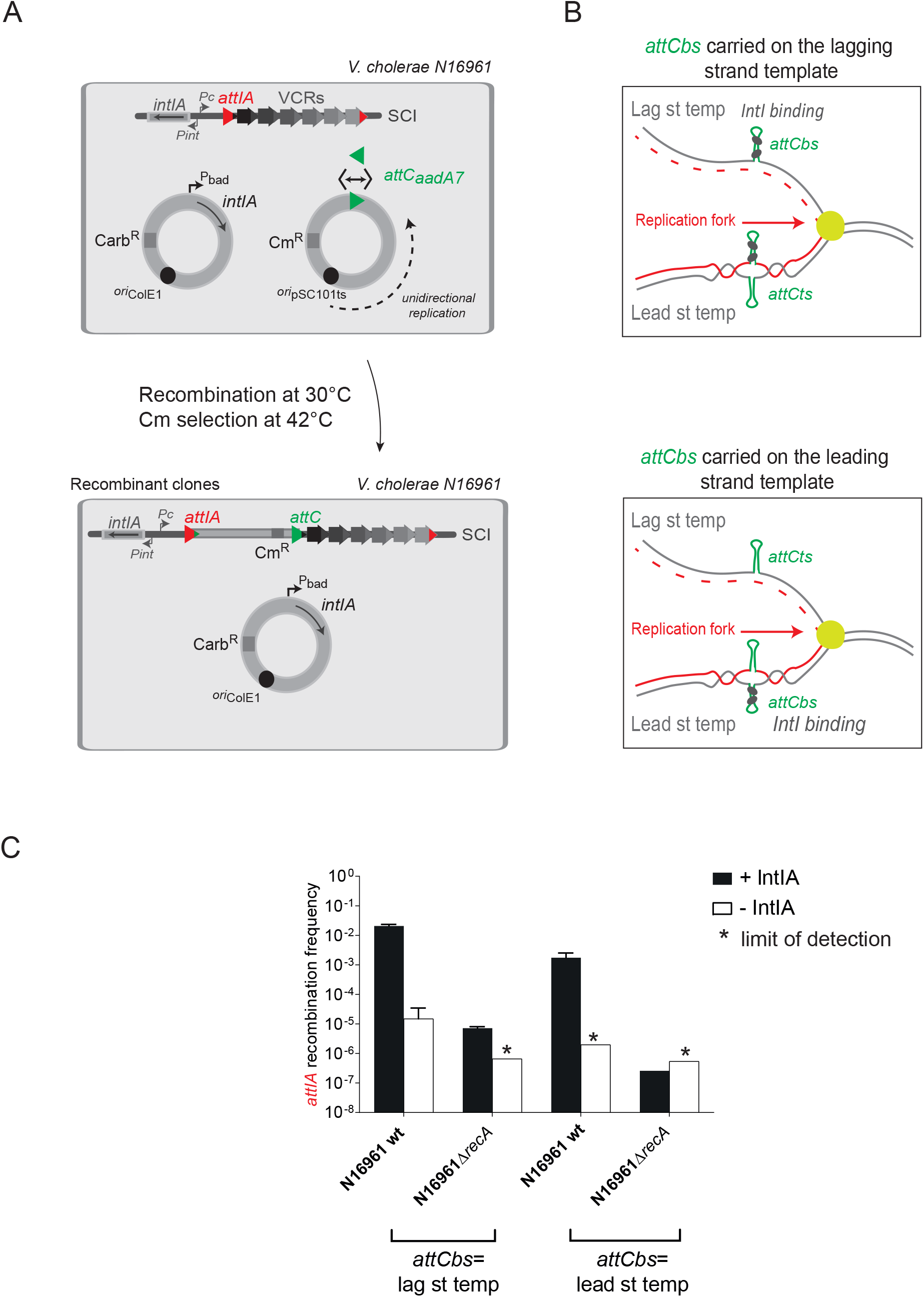
Effect of the RecA protein on *attC* × *attI* reaction when *attC* sites are carried by a replicative vector in *V. cholerae.* (A) Experimental setup of the replicative assay. *V. cholerae* N16961 strains transformed with both pBAD18 IntIA expressing vector and the unidirectional-replicative vector, pTSC29::*attC*_*aadA7*_ were used. *attCaadA7* sites (green triangles) were cloned in both orientations (double arrow) in the pTSC29 vector. Since pTSC29 has a thermosensitive origin of replication, recombination reaction is performed at 30°C and, to evaluate the recombination frequency, recombinant clones selection was performed on Cm containing plates at 42°C (see also Results and Material and methods). The *attIA* site on the *V. cholerae* SCI is represented by a red triangle. (B) Folding of *attC*_*aadA7*_ site on replicative vector. Replicated and template strands are coloured in red and grey respectively and grey circle represent IntIA monomers. bs: bottom strand; ts: top strand; Lag st temp: Lagging strand template; Lead st temp: Leading strand template. (C) Frequency of insertion of the pTSC29::*attC*_*aadA7*_ unidirectional-replicative vector into the *attIA* site. The recombination rates were calculated in N16961 *V. cholerae* wt and in the corresponding *recA* mutant strains (*ΔrecAΔ*). Under plots, the orientation of *attC*_*aadA7*_ bs on lagging or leading strand template (lag st temp or lead st temp) are indicated. Results correspond to recombination frequencies that were normalized after analysis of PCR reactions (Material and Methods). +IntI: recipient strains transformed with the pBAD18 integrase expressing vector; -IntI: control strains transformed with the empty pBAD18 vector. * correspond to the limits of detection. Values represent the mean of at least three independent experiments and error bars correspond to average deviations from the mean.

## DISCUSSION

### *Incoming integron cassettes are efficiently inserted in the attIA site of the* Vibrio cholerae integron

Until now, integron cassette movements in natural integrons, mediated by the endogenous integrase, have never been experimentally demonstrated. Indeed, exchange of gene cassettes between bacterial species (Domingues et al., 2012) and gene cassette movements in the *V. cholerae* SCI (Baharoglu et al., 2010) have been detected but at very low frequencies and related to homologous recombination events. Here, for the first time, we worked with the entire endogenous SCI of the *Vibrio cholerae* pathogen strain and we succeed to visualize cassette recombination events dependent of the sole endogenous integron integrase. We used, to deliver the integron cassettes, both conjugation and natural transformation processes. These two frequent mechanisms perfectly mimic the natural conditions in which cassettes can be delivered through horizontal gene transfer in the *Vibrio cholerae* strains. When delivering cassettes by conjugation assay, we detected high rate of cassette insertion (~10^−2^) in conditions where IntIA is overexpressed. However, we were also able to detect a significant level of cassette insertion (>10^−6^) catalyzed by the sole endogenous integrase expression. We also detect cassette insertion events catalyzed by the endogenous integrase by delivering cassette during natural transformation. The entry of single-stranded cassettes by natural gene transfer processes activates the SOS response sufficiently to express the endogenous integrase leading to the proper insertion of these incoming cassettes. We therefore reproduce the whole natural integron pathway in which incoming single-stranded integron cassettes by HGT processes activates their proper insertion in the integron platform (Figure 8).

**Figure 8:**
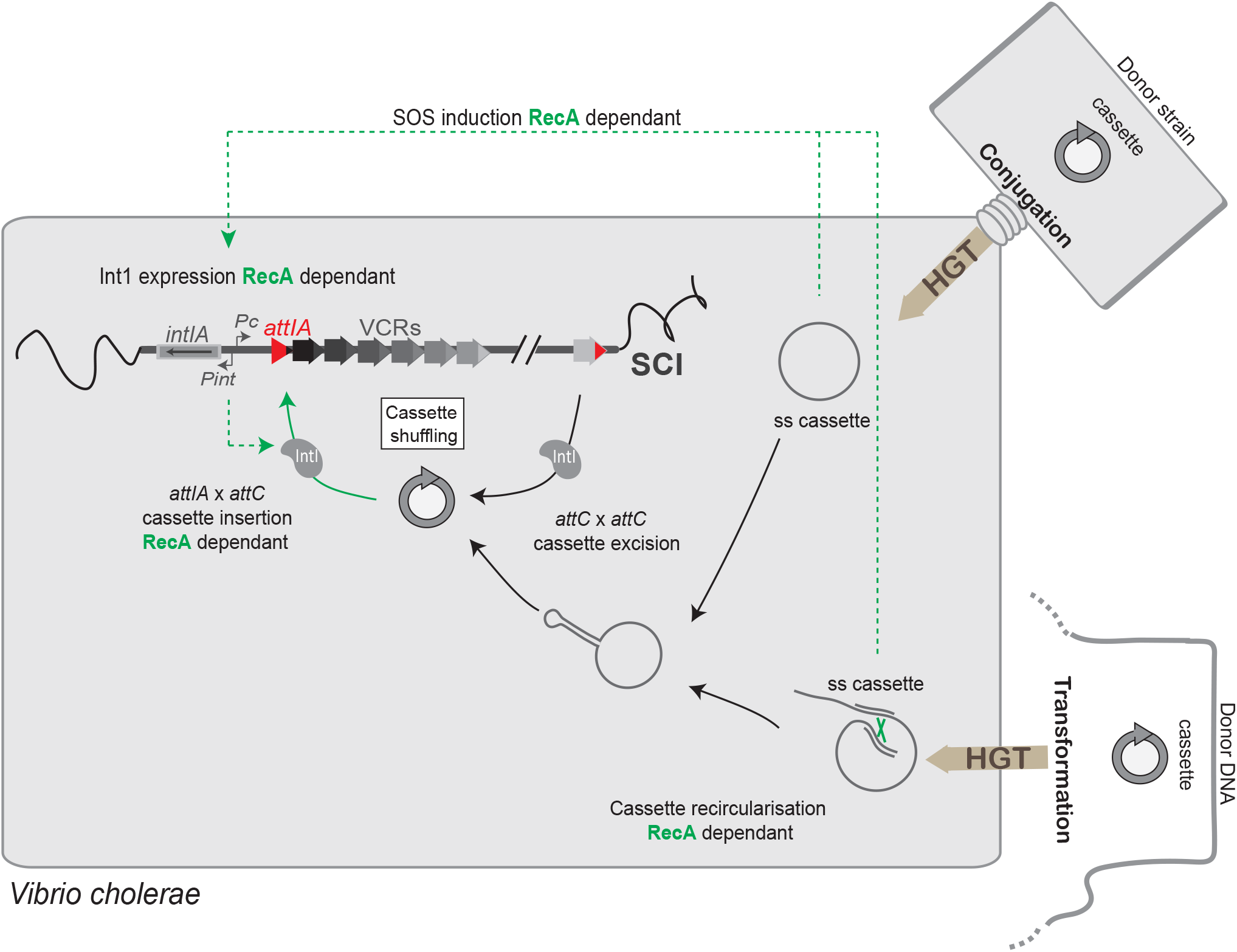
Snap shot of SCI recombination during horizontal gene transfer *in Vibrio cholerae*. The SCI activity is represented and its connections with bacterial physiology. The steps, which involve the RecA protein, are indicated in green. HGT: horizontal gene transfer; ss: single-stranded; VCR: *Vibrio cholerae* Repeat sequences; SCI: sedentary chromosomal integron.

Interestingly, by analyzing the insertion events, we demonstrated that despite the presence of 180 VCR sites in the large array of cassettes, the *attIA* site constitutes the major recruitment site for cassette insertion in the *V. cholerae* SCI. This recruitment, in the first position of the cassette array, ensures to the newly inserted cassettes to be efficiently expressed. This meaning that, even if the frequency of cassette insertion seems low when mediated by the endogenous integrase (~10^−6^), these events can be selected and evolutionary fixed if cassette expression leads to an advantageous new function, which ensures an adaptive response to one environmental selective pressure. Thus, the knowledge of the precise cassette functions in SCI would allow to trace back the environmental conditions of cell growth and the evolutionary history of SCI-containing cells.

### *RecA*_Vch_ *is critical for cassette insertion in the* **attIA** *site of the* Vibrio cholerae SCI

Previous studies have shown how integrons are host-cell integrated systems. For instance, the folding of single-stranded *attC* substrates depends on several cellular processes including conjugation, replication or DNA topology (Loot et al., 2010; Loot et al., 2017). *attC* site folding can also be modulated by the binding of the SSB host protein, which, by hampering *attC* hairpin folding *in vivo* in the absence of IntI, would play an important role in maintaining the *attC* integrity and its recombinogenic functionality (Loot et al., 2014). It has also been shown that the pathway that allows the resolution of aHJ formed during *attI* × *attC* reaction is linked to a host-replicative process (Loot et al., 2012). Moreover, integrase expression is also extensively regulated by environmental stresses, thus connecting cassette recombination to the host-cell environment. Both integrase and cassette expressions, ensured respectively by P_int_ and P_C_, are regulated by catabolite repression in *V. cholerae* SCI (Baharoglu et al., 2012; Krin et al., 2014). Expression of the MI class 1 integrase is in part regulated by the stringent response in biofilms (Strugeon et al., 2016). The most relevant example among these regulatory pathways, is undoubtedly the induction of integrase expression (of class 1 MI and *V. cholerae* SCI) during the SOS response. Such regulation is ensured by the binding of the LexA repressor in the promoter region of P_int_ (Cambray et al., 2011; Guerin et al., 2009). Integrase expression like for all the other genes of the SOS regulon (more than 40 SOS genes), is unclenched by the autocatalytic cleavage of LexA, a process induced by the binding of RecA on single-stranded DNA constituting nucleofilament. This triggered SOS response is a global response to DNA damage ensuring a state of high-activity DNA repair. In this study, we investigated the role of the RecA protein in another regulatory point of the integron system, the cassette recombination. We choose to study this protein for two reasons. First, because of its ssDNA binding properties, since we know that cassette recombination involves one of the recombination sites, the *attC* site, in a single-stranded DNA form. Second, because efficient cassette recombination could be dependent on one or several proteins belonging to the SOS regulon and whose expression is induced in presence of RecA. We found that the RecA_*Vch*_ protein is critical for the *attIA* × *attC* reaction in the SCI of *V. cholerae*. On contrary, when we tested the effect of RecA_*Vch*_ on cassette insertions in VCR sites of the SCI, we found that these *attC* × VCR reactions were independent of the RecA_*Vch*_ protein. We also confirmed the absence of RecA effect on *attC* × *attC* recombination by testing cassette insertion in an *attC*_*aadA1*_ site-carrying plasmid. Interestingly, these results show that the RecA effect depend on the nature of *att* sites (*attI* or *attC*), which are recombined. Altogether, our results indicate that the RecA protein is a host factor specifically implicated in the cassette recruitment in the *attIA* site of the *V. cholerae* SCI during environmental stress conditions. Finally, RecA protein acts at several steps during integron recombination process (Figure 8). First, as said above, when antibiotic is used or when single-stranded cassette are released in cell by HGT, RecA favors the expression of the SOS-dependent integrase gene (Baharoglu and Mazel, 2011; Guerin et al., 2009). Second, since *recA* gene itself also belongs to the SOS regulon (Courcelle et al., 2001; Little et al., 1981), its expression is up-regulated (ten-fold within minutes (Courcelle et al., 2001; Renzette et al., 2005)). This release of RecA protein would allow RecA to be available to trigger its biological functions, upon which its stimulating effect on cassette recruitment at *attIA* SCI (Figure 8).

### *Model of RecA*V_ch_ *mechanism of action*

RecA activates expression of a large number of genes during the SOS response (Krin et al., 2018). The mechanism of action of RecA_*Vch*_ on integron recombination could thus be the consequence of an indirect effect. Using a *V. cholerae* mutant strain *lexAind-* in which the SOS response is constitutively repressed, we showed that SOS response abrogation did not affect recombination rate and therefore that the observed RecA_*Vch*_ effect on *attIA* × *attC* recombination does not involve any other protein belonging to the SOS regulon. Besides, we know that the *E. coli* RecA protein differs from the *V. cholerae* RecA protein in the extreme C-terminal part and that this part is known to modulate interactions with regulatory proteins (Figure S1) (Cox, 2007b). We decided to study the effect of the RecA_*Ec*_ protein in *attIA* × *attC* recombination mediated by IntIA. We performed complementation of the *V. cholerae ΔrecA* mutant by the *recA*_*Ec*_ gene. We observed that the RecA_*Ec*_ protein fully restores the recombination activity, meaning that both RecA proteins may insure cassette insertion in the *attIA* site.

Contrary to the *attC* × *attIA* reaction, we did not observe any influence of RecA_*Vch*_ on *attC* × *attC* reaction mediated by IntIA in *V. cholerae*, while both reactions require ss *attC* site on double-stranded DNA molecule, we still observed an important effect of the RecA_*Vch*_ protein. Then, whatever the way to deliver *attC* sites, i.e. single-stranded (conjugation and replicative assays) or double-stranded (replicative assay), we observed an important effect of the RecA_*Vch*_ protein in *attIA* × *attC* recombination. Altogether, these results show that observed RecA_*Vch*_ effect is not linked to the regulation of the *attC* sites folding. We therefore suppose that the *attIA* × *attC* synaptic complex formed during the recombination could impede the aHJ resolution by blocking replication fork progression. The effect of RecA would be to bind the ss *attC* sites and help to maintain the *attC* site in a non-folded single-stranded form destabilizing the synaptic complex and favoring the replicative resolution step. Note that we did not observe this RecA_*Vch*_ effect on *attC* × *attC* recombination suggesting a differential configuration between both *attC* × *attC* and *attIA* × *attC* complexes. Indeed, synapses formed during both reactions are known to be different since involving respectively two flexible bs *attC* sites and, a flexible bs *attC* and a stiffer ds *attI* (Demarre et al., 2007). Such differences would explain that the synapse architecture for the *attIA* × *attC* reaction versus *attC* × *attC* may potentially requires the assistance of additional accessory proteins. Furthermore, the *attC* × *attC* reaction can be catalyzed by IntIA in *E. coli* while, productive *attIA* × *attC* reactions are almost undetectable in this host (Biskri et al., 2005).

### Host factor recruitment differs between Mobile and Sedentary Chromosomal Integrons

Biskri *et al.* previously observed the integrase of the SCI of *V. cholerae* recombines 2 000 fold less efficiently during *attIA* × *attC* reaction, when expressed in a heterologous host such as *E. coli* (Biskri et al., 2005). This observation suggests that IntIA requires host factors that are absent or too divergent in *E. coli* to carry out this reaction (Biskri et al., 2005). Here, we confirmed this result and failed at recovering *attIA* × *attC* recombination by expressing RecA_*Vch*_ in *E. coli*, suggesting that the RecA_*Vch*_ protein is not the missing factor in *E. coli* impeding this reaction to efficiently occur. This is not surprising since, even though they show some slight variations in the C terminal part, RecA_*Ec*_ and RecA_*Vch*_ present a very high level of identity (80%, Figure S2). Besides, this is in accordance with the fact that RecA_*Ec*_ was able to functionally complement the absence of RecA_*Vch*_ to achieve this reaction in *V. cholerae* (Figure 3A). Therefore, we hypothesize that other specific *V. cholerae* host factors are missing in *E. coli*.

In contrast, RecA is not involved in any reactions mediated by IntI1 (neither in *E. coli* or *V. cholerae*) and all these reactions are efficient in *V. cholerae*. Therefore, while SCI cassette recombination activity seems restricted to a given host, MI recombination seems efficient in different species suggesting a host factor recruitment more stringent for SCIs compare to MIs. This is consistent with the observed widespread MIs dissemination among bacterial species. The evolutionary success of class 1 MIs clearly reflects the fact that they are functional in a wide range of bacterial hosts. This could result from a co-evolution between the integrase IntI1 and its *attI1* site to allow that cassette recombination takes place in absence of specific accessory proteins. Both IntI and *attI* level of divergence, 45% between IntIA and IntI1 (Demarre et al., 2007) and the lack of identity between *attIA* and *attI1* sites, likely reflect their functioning differences. Indeed, *attI1* site harbors supplementary IntI1 binding sites, the direct repeats (DRs, Figure S3), that constitute accessory sequences to which the integrase is able to bind (Gravel et al., 1998). These DRs favor *attI1* × *attC* reaction (Partridge et al., 2000). However, such sequences are not common features to the majority of *attI* sites (Nield et al., 2001) and, for instance, the *attIA* site of the *V. cholerae* SCI does not harbor any DRs (Figure S3). Upon mobilization of integrons, *attI* sites could have evolved and acquired such motifs (e.g. DRs), to replace trans-acting host factors necessary for the *attI1* × *attC* reactions. For instance, DRs of *attI1* site acting as topological filters (Partridge et al., 2000) could therefore replace host factors that regulate supercoiling.

### Natural transformation represents an efficient pathway for SCI cassette recruitment

HGT contributes to the emergence of pathogens and the spread of virulence factors, and also, enables many disease-causing bacteria to rapidly evolve in response to environmental pressures such as antibiotic use (Blokesch, 2016; de la Cruz and Davies, 2000; Frost et al., 2005; Gogarten et al., 2002; Ochman et al., 2005; Prudhomme et al., 2006; Waldor and Mekalanos, 1996). Among HGT processes, natural competence is able to mediate the absorption and exchange of free DNA when sufficient homology is present between the incoming DNA and the bacterial genome (Chen and Dubnau, 2004; Griffith, 1928). Here, we demonstrated that acquisition of material during natural transformation can be also mediated by homology independent DNA mechanism, notably by exchanging and recruiting gene cassettes inside integrons in an integrase dependent way. We used the causative agent of the diarrheal disease cholera, *Vibrio cholerae*, responsible for seven major pandemics since 1817, the latter of which is still ongoing (Lippi et al., 2016). Cassette recruitments occur directly in the *attIA* site of the SCI contained in *V. cholerae* where they are expressed and selected if procuring an advantage to the *Vibrio cholerae* strain. The natural bacterial competence has already been demonstrated for at least 80 species of bacteria but remains little explored and is probably underestimated (Johnsborg et al., 2007; Johnston et al., 2014). Notably, natural transformation is largely widespread among *Vibrionaceae* where SCIs are mainly found and constitute very plastic regions (Le Roux and Blokesch, 2018; Matthey et al., 2019). Here, we validated a new pathway for integron cassette recruitment in SCIs triggered by natural transformation probably involved in adaptation and making integrons important motors of evolution in *Vibrionaceae* species.

## SUPPLEMENTARY DATA

Table S1: Bacterial strains used in this study

Table S2: Plasmids used in this study

Table S3: Primers used in this study

Figure S1: Electrophoretic analysis of reactional substrates

Figure S2: Alignment of the RecA_*Ec*_ and RecA_*Vch*_ protein sequences

Figure S3: *attI* site structures

## ACKNOWLEDGEMENT

We would like to thank Evelyne Krin for providing the N16961 *recA* mutant. We also thank all the lab members for helpful discussion. We also thank Sandra Arroyo-Beck and Marcos Manero for their experimental help.

