## Supplementary material for "Cassette recruitment in the chromosomal Integron of *Vibrio cholerae*": Supplemn

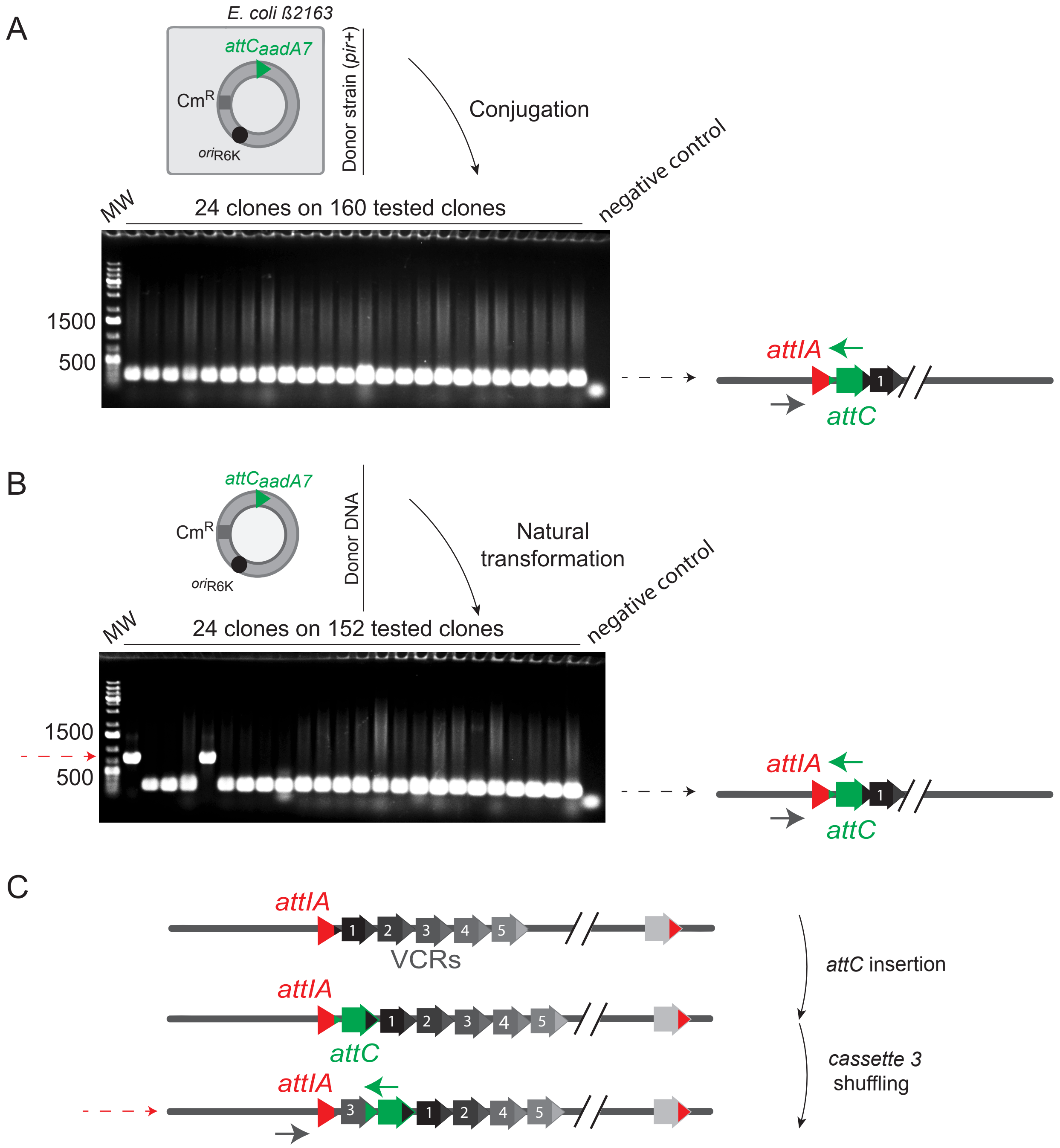


**Figure S1**: **Analysis of recombined clones**

A and B. Electrophoresis analysis of PCR products obtained on recombined clones after conjugation (A) and natural transformation assays (B). Reactional products and used primers for PCR (grey and green arrows) are shown. The *attCaadA7* site carried by the suicide vector is represented by a green triangle. MW: Molecular Weight Marker DNA

(C) Schemes showing the successing events (*attCaadA7* cassette insertion followed by cassette 3 shuffling) leading to reactional products showed by the dashed red arrow in B .

The *attIA* site on the *V. cholerae* SCI is represented by a red triangle and VCR sites by arrows.

```
RecAEc     1 MAIDENKQKALAAALGQIEKQFGKGSIMRLGEDRSMDVETISTGSLSLDIALGAGGLPMG  60
               +DENKQKALAAALGQIEKQFGKGSIMRLG++R+MDVETISTGSLSLDIALGAGGLPMG
RecAVch    1 --MDENKQKALAAALGQIEKQFGKGSIMRLGDNRAMDVETISTGSLSLDIALGAGGLPMG  58

RecAEc    61 RIVEIYGPESSGKTTLTLQVIAAAQREGKTCAFIDAEHALDPIYARKLGVDIDNLLCSQP 120
             RIVEI+GPESSGKTTLTL++IAAAQREGKTCAFIDAEHALDP+YA+KLGV+ID LL SQP
RecAVch   59 RIVEIFGPESSGKTTLTLELIAAAQREGKTCAFIDAEHALDPVYAKKLGVNIDELLVSQP 118

RecAEc   121 DTGEQALEICDALARSGAVDVIVVDSVAALTPKAEIEGEIGDSHMGLAARMMSQAMRKLA 180
             DTGEQALEICDALARSGAVDVIVVDSVAALTPKAEIEGE+GDSHMGL ARM+SQAMRKL
RecAVch  119 DTGEQALEICDALARSGAVDVIVVDSVAALTPKAEIEGEMGDSHMGLQARMLSQAMRKLT 178

RecAEc   181 GNLKQSNTLLIFINQIRMKIGVMFGNPETTTGGNALKFYASVRLDIRRIGAVKEGENVVG 240
             GNLKQSN + IFINQIRMKIGVMFGNPETTTGGNALKFYASVRLDIRR GA+KEGE VVG
RecAVch  179 GNLKQSNCMCIFINQIRMKIGVMFGNPETTTGGNALKFYASVRLDIRRTGAIKEGEEVVG 238

RecAEc   241 SETRVKVVKNKIAAPFKQAEFQILYGEGINFYGELVDLGVKEKLIEKAGAWYSYKGEKIG 300
             +ETR+KVVKNKIAAPFK+A  QI+YG+G N  GEL+DLGVK K++EK+GAWYSY G+KIG
RecAVch  239 NETRIKVVKNKIAAPFKEANTQIMYGQGFNREGELIDLGVKHKMVEKSGAWYSYNGDKIG 298

RecAEc   301 QGKANATAWLKDNPETAKEIEKKVRELLLSNPNS---TPDFSVDDSEGVAETNEDF 353
             QGKANA  +LK+NPE AK ++KK+RE+LL+  N        S  D        E+F
RecAVch  299 QGKANACKYLKENPEIAKTLDKKLREMLLNPENMQLIAETSSAADDVEFGAVPEEF 354
```

**Figure S2**: *Alignment of the RecAEc and RecAVch protein sequences*

The alignment was made on Geneious. Identical amino acids are listed between the two sequences as well as functionally conservative substitutions (shown by +).

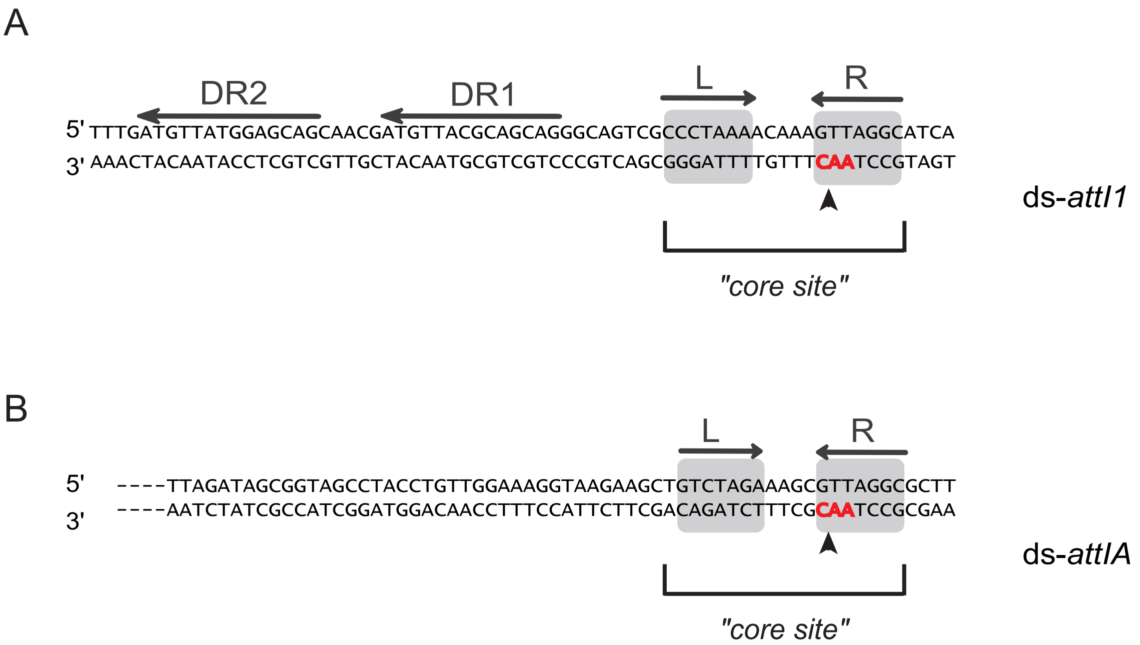


**Figure S3: Schematic representation of the double-stranded *attI* recombination sites**

The sequences of the double-stranded (ds) *attI* sites of the class 1 MI (A) and of the SCI of *Vibrio cholerae* platforms (B) are represented, respectively *attI1* and *attIA*. Grey boxes indicate the R (right) and L (left) integrase binding sites (core site). The 5'-AAC-3' triplet, where the cleavage takes place is highlighted in red and the precise cleavage point is indicated by a black arrowhead. The Direct Repeats (DR1 and DR2) of the *attI1* site are shown by black arrows.

*Table S1: Bacterial strains used in this study*

| Strain number | Relevant genotypes or description | References |
|---|---|---|
| | **Basic *V. cholerae* strains** | |
| 7805 | N16961 biovar El Tor | Laboratory collection |
| 8637 | N16961 *hapR*+ [$Str^R$ $Gm^R$] | Laboratory collection |
| 9156 | N16961 *hapR*+ *ΔintIA* [$Str^R$ $Gm^R$] | (Le Roux et al., 2007) |
| B306 | 7805 *ΔrecA* | This study, deletion of the *recA* gene (VC0543) was performed by allelic exchange by delivering the suicide conjugative plasmid pB203 in the 7805 strain. |
| H858 | 8637 *lexAind*- (A91D) [$Str^R$ $Gm^R$] | This study, point mutation in *lexA* gene was introduced by allelic exchange by delivering the suicide conjugative plasmid p6780 (Guerin et al., 2009) in the 8637 strain. |
| L438 | B306 *att*Tn7::$P_{LAC}$-*recA*$_{Ec}$ | This study, $P_{LAC}$-recA$_{Ec}$ was inserted into the *att*Tn7 locus following the previously described protocol (de Lemos Martins et al., 2018) and by delivering by conjugation the pL310 shuttle vector into the B306 strain |
| L435-37 | B306 *att*Tn7::$P_{LAC}$-*recA*$_{Vch}$ | This study, $P_{LAC}$-*recA*$_{Vch}$ was inserted into the *att*Tn7 locus following the previously described protocol (de Lemos Martins et al., 2018) and by delivering by conjugation the pL312 shuttle vector into the B306 strain. |
| K779-K781 | 7805 *ΔattIA* | This study, *attIA* site was deleted by allelic exchange by delivering the suicide conjugative plasmid pK590 in the 7805 strain. |
| K782-K784 | B306 *ΔattIA* | This study, *attIA* site was deleted by allelic exchange by delivering the suicide conjugative |

| | | |
|---|---|---|
| | | plasmid pK590 in the B306 strain that was previously transformed with the pAM::*recA*$_{Ec}$ vector. |
| K756-K757-K955 | 7805 *ΔattIA::attI1* | This study, *attIA* site was replaced by allelic exchange by delivering the suicide conjugative plasmid pK584 in the 7805 strain. |
| K956-K958 | B306 *ΔattIA::attI1* | This study, *attIA* site was replaced by allelic exchange by delivering the suicide conjugative plasmid pK584 in the B306 strain that was previously transformed with the pAM::*recA*$_{Ec}$ vector. |
| | **Basic *E. coli* strains** | |
| 8195 | DH5α, (F-) *supE44 ΔlacU169* (ϕ80lacZΔM15) *ΔargF hsdR17 recA1 endA1 gyrA96 thi-1 relA1* | Laboratory Collection |
| 4196 | β2163, (F-) RP4-2-Tc::Mu *ΔdapA*::(*erm-pir*) [Km$^R$ Em$^R$] | (Demarre et al., 2005) |
| 8725 | π3813, *lacIq thi-1 supE44 endA1 recA1 hsdR17 gyrA462 zei-298::Tn10 ΔthyA*::(*erm-pir116*) [Em$^R$] | (Le Roux et al., 2007) |
| 8726 | β3914, β2163 *gyrA462 zei-298::Tn10* [Km$^R$ Em$^R$] | (Le Roux et al., 2007) |
| C349 | MG1655 *E. coli* K12 wt | Laboratory collection |
| 4826 | MG1655 *recA269::Tn10* [Tc$^R$] | RG Lloyd |
| L805-L806 | 4826 *att*Tn7::P$_{LAC}$-*recA*$_{Vch}$ [Tc$^R$] | This study, P$_{LAC}$-recA$_{Vch}$ fragment was inserted into the *att*Tn7 locus following the previously described protocol (de Lemos Martins et al., 2018) by delivering the pL312 shuttle vector into the 4826 strain |

| | **Transformed strains of *V. cholerae* used in suicide conjugation assay** | |
|---|---|---|
| J554-J556 | N16961 pJ502 | This Study |
| J529-J531 | N16961 pJ504 | This Study |
| J566-J568 | N16961 *hapR*+ pJ502 | This Study |
| J541-J543 | N16961 *hapR*+ pJ504 | This Study |
| o150-o152 | N16961 *hapR*+ *ΔintIA* pJ502 | This Study |
| o153-o155 | N16961 *hapR*+ *ΔintIA* pJ504 | This Study |
| J557-J559 | N16961*ΔrecA* pJ502 | This Study |
| J532-J534 | N16961*ΔrecA* pJ504 | This Study |
| J569-J571 | N16961 *hapR*+ *lexAind*- pJ502 | This Study |
| J544-J546 | N16961 *hapR*+ *lexAind*- pJ504 | This Study |
| L453 | N16961*ΔrecA att*Tn7::$P_{LAC}$-$recA_{Ec}$ pJ502 | This Study |
| L447 | N16961*ΔrecA att*Tn7::$P_{LAC}$-$recA_{Ec}$ pJ504 | This Study |
| L449-L451 | N16961*ΔrecA att*Tn7::$P_{LAC}$-$recA_{Vch}$ pJ502 | This Study |
| L443-L445 | N16961*ΔrecA att*Tn7::$P_{LAC}$-$recA_{Vch}$ pJ504 | This Study |
| M933 | N16961*ΔattIA* pJ502 pM923 | This Study |
| M927 | N16961*ΔattIA* pJ504 pM923 | This Study |
| N457-N459 | N16961*ΔrecA ΔattIA* pJ502 pM923 | This Study |
| N454-N456 | N16961*ΔrecA ΔattIA* pJ504 pM923 | This Study |
| K870-K872 | N16961*ΔattIA* pJ502 | This Study |
| K864-K866 | N16961*ΔattIA* pJ504 | This Study |
| K873-K875 | N16961*ΔrecA ΔattIA* pJ502 | This Study |
| K867-K869 | N16961*ΔrecA ΔattIA* pJ504 | This Study |
| L326-L328 | N16961*ΔattIA* p2597 | This Study |
| L329-L331 | N16961*ΔattIA* p8532 | This Study |
| L338-L340 | N16961*ΔrecA ΔattIA* p2597 | This Study |
| L341-L343 | N16961*ΔrecA ΔattIA* p8532 | This Study |
| K976-K978 | N16961*ΔattIA::attI1* pL290 | This Study |
| K970-K972 | N16961*ΔattIA::attI1* pL294 | This Study |

| | | |
|---|---|---|
| K979-K981 | N16961*ΔrecA ΔattIA::attI1* pL290 | This Study |
| K973-K975 | N16961*ΔrecA ΔattIA::attI1* pL294 | This Study |
| L581-L583 | N16961*ΔattIA::attI1* p929 pL290 | This Study |
| L584-L586 | N16961*ΔattIA::attI1* p929 pL294 | This Study |
| L713-L715 | N16961*ΔrecA ΔattIA* p929 pL290 | This Study |
| L716-L718 | N16961*ΔrecA ΔattIA* p929 pL294 | This Study |
| | **Transformed strains of *V. cholerae* used in suicide natural transformation assay** | |
| N389-N390 | N16961 *hapR*+ pM889 | This Study |
| N387-N388 | N16961 *hapR*+ pN346 | This Study |
| N535-N537 | N16961 *hapR*+ *ΔintIA* pM889 | This Study |
| N532-N534 | N16961 *hapR*+ *ΔintIA* pN346 | This Study |

| | **Transformed strains of *E. coli* in suicide conjugation assay** | |
|---|---|---|
| L120 | β2163 pD060 | This study |
| L385-L387 | MG1655 p929 pL290 | This study |
| J733-J735 | MG1655 p929 pL294 | This study |
| L388-L390 | MG1655 p1105 pL290 | This study |
| K689-K690 | MG1655 p1105 pL294 | This study |
| N369-N371 | MG1655 pM923 pJ502 | This study |
| N372-N374 | MG1655 pM923 pJ504 | This study |
| K686-K688 | MG1655 p1105 pJ502 | This study |
| K683-K685 | MG1655 p1105 pJ504 | This study |
| L391-L393 | MG1655 *recA269::Tn10* p929 pL290 | This study |
| J990-J991 | MG1655 *recA269::Tn10* p929 pL294 | This study |
| L394-L396 | MG1655 *recA269::Tn10* p1105 pL290 | This study |
| K698-K699-J992 | MG1655 *recA269::Tn10* p1105 pL294 | This study |

| | | |
|---|---|---|
| N375-N377 | MG1655 *recA269::Tn10* pM923 pJ502 | This study |
| N378-N380 | MG1655 *recA269::Tn10* pM923 pJ504 | This study |
| K695-K697 | MG1655 *recA269::Tn10* p1105 pJ502 | This study |
| K692-K694 | MG1655 *recA269::Tn10* p1105 pJ504 | This study |
| N381-N383 | MG1655 *recA269::Tn10- att*Tn7::$P_{LAC}$-*recA*$_{Vch}$ pM923 pJ502 | This study |
| N384-N386 | MG1655 *recA269::Tn10- att*Tn7::$P_{LAC}$-*recA*$_{Vch}$ pM923 pJ504 | This study |

| | **Transformed strains of *V. cholerae* used in recombination assay with unidirectional-replicative substrate** | |
|---|---|---|
| K160-K162 | N16961 p979 p7523 | This study |
| K169-K171 | N16961 p979 p7546 | This study |
| K172-K174 | N16961 p995 p7523 | This study |
| K181-K183 | N16961 p995 p7546 | This study |
| K184-K186 | N16961*ΔrecA* p979 p7523 | This study |
| K193-K195 | N16961*ΔrecA* p979 p7546 | This study |
| K196-K198 | N16961*ΔrecA* p995 p7523 | This study |
| K205-K207 | N16961*ΔrecA* p995 p7546 | This study |

*Table S2: Plasmids used in this study*

| Plasmid number | Plasmid description | Relevant properties and construction |
|---|---|---|
| p452 | pSU18Δ::*VCR*$_{128}$ | *ori*p15A; [Cm$^R$] (Biskri et al., 2005) |
| p741 | pSU38Δ | *ori*p15A; [Km$^R$] (Biskri et al., 2005) |
| p755 | pSU38Δ::*attIA* | *ori*p15A; [Km$^R$] (Biskri et al., 2005) |
| p929 | pSU38Δ::*attI1* | *ori*p15A; [Km$^R$] (Biskri et al., 2005) |

| | | |
|---|---|---|
| p1105 | pSU38Δ::VCR$_{VCA0441}$ | *ori*p15A; [Km$^R$] EcoRI/BamHI fragment (VCR) from p452 cloned in EcoRI/BamHI digested p741 (Biskri, unpublished results) |
| p7523 | pTSC29::*attC*$_{aadA7}$ (ori-) ("lead") | *oripSC101*ts; [Cm$^R$] (Loot et al., 2010) |
| p7546 | pTSC29::*attC*$_{aadA7}$ (ori+) ("lag") | *oripSC101*ts; [Cm$^R$] (Loot et al., 2010) |
| p979 | pBAD18 | *ori*ColE1; [Carb$^R$] (Guzman et al., 1995) |
| p995 | pBAD18::*intIA*$_{Vch}$ | *ori*ColE1; [Carb$^R$] (Biskri et al., 2005) |
| p3938 | pBAD18::*intI1* | *ori*ColE1; [Carb$^R$] (Demarre et al., 2007) |
| p2597 | pBAD43 | *oripSC101*; [Sp$^R$] (Guzman et al., 1995) |
| p8532 | pBAD43::*intIA*$_{Vch}$ | *oripSC101*; [Sp$^R$] (Biskri, unpublished) |
| pG906 | pBAD43::*intI1* | *oripSC101*; [Sp$^R$] EcoRI/HindIII digested fragment (*intI1*) from p3938 cloned in EcoRI/HindIII digested p2597 |
| pJ502 | pBAD43*ΔattC*$_{aadA1}$ | *oripSC101*; [Sp$^R$] amplification by inverse PCR of p2597 with primers 5640 and 5641 |
| pJ504 | pBAD43 *ΔattC*$_{aadA1}$::*intIA*$_{Vch}$ | *oripSC101*; [Sp$^R$] amplification by inverse PCR of p8532 with primers 5640 and 5641 |
| pL290 | pBAD43::*aadA7* | *oripSC101*; [Sp$^R$] The fragment P$_{cat}$-*aadA7* was amplified from pSW25T vector (Demarre et al., 2005) with primers 5234 and 5235. pJ502 vector was amplified by inverse PCR with 5914 and 5915 primers. Assembly of these two fragments was achieved by performing Gibson Assembly. |
| pL294 | pBAD43::*aadA7-intI1* | *oripSC101*; [Sp$^R$] The same strategy was used as for the construction of pL290 but using pG906 as template for vector amplification with 5914 and 5915 primers. |
| pM783 | pSU38Δ::*intIA-attIA* | *ori*p15A; [Km$^R$] The *intIA-attIA* fragment was amplified from the 7805 strain using o6176 and o6177 primers. This fragment was digested with |

| | | |
|---|---|---|
| | | EcoRI and BamHI restriction enzymes and ligated into the EcoRI and BamHI digested p755 vector. |
| pM923 | pSU38Δ::*intIAY302F-attIA* | *ori*p15A; [$Km^R$] Inverse PCR was performed using the pM783 plasmid and, o6184 and o6185 primers. |
| pE639 | pLC10 | *oripSC101ts*; *oripSC101*; [$Sp^R$] Laboratory of David Bikard, Institut Pasteur |
| p7301 | pCY579 (pAM::$recA_{Ec}$) | *oripSC101*; [$Carb^R$] (Cronan, 2003) |
| pM889 | pBAD43$\Delta attC_{aadA1}$ -$P_{tet}$ | *oripSC101*; [$Sp^R$] The $P_{tet}$ promoter fragment was amplified from pE639 vector using 6178 and 6182 primers. pJ502 vector was amplified by inverse PCR with 6181 and 6183 primers. Assembly of these two fragments was achieved by performing Gibson Assembly. |
| pN346 | pBAD43$\Delta attC_{aadA1}$-$P_{tet}$-*intIA* | *oripSC101*; [$Sp^R$] The $P_{tet}$ promoter fragment was amplified from pE639 vector using 6178 and 6230 primers. pJ504 vector was amplified by inverse PCR with 6181 and 6229 primers. Assembly of these two fragments was achieved by performing Gibson Assembly. |
| pD060 | pSW23T::$attC_{aadA7}$ (bs) | $oriV_{R6K\gamma}$, $oriT_{RP4}$; [$Cm^R$] (Nivina et al., 2016) |
| p7848 (pMP7) | Suicide conjugative plasmid used for allelic exchange pSW23T-*araC* $P_{BAD}$-*ccdB* | $oriV_{R6K\gamma}$, $oriT_{RP4}$; [$Cm^R$] (Val et al., 2012) |
| p6780 | Suicide conjugative plasmid used for replacement of *lexAind-* allele | $oriV_{R6K\gamma}$, $oriT_{RP4}$; [$Cm^R$] (Guerin et al., 2009) |
| pB203 | Suicide conjugative plasmid used for deletion of $recA_{Vch}$ gene (VC0543) | $oriV_{R6K\gamma}$, $oriT_{RP4}$; [$Cm^R$] homology regions upstream and downstream of the VC0543 gene were amplified with VC0542A and RECAB and with RECAC and RECAD primers respectively. For both PCR the N16961 genomic DNA was used as a template. |

| | | |
|---|---|---|
| | | These two fragments were assembled by PCR using VC0542A and RECAD primers. This combined fragment was digested by EcoRI and cloned into EcoRI digested p7848 vector. (Krin, unpublished) |
| pK584 | Suicide conjugative plasmid used for deletion of *attIA* and replacement by *attI1* | $oriV_{R6K\gamma}$, $oriT_{RP4}$; [$Cm^R$] amplification of fragments upstream and downstream of *attIA* were performed using 5738 and 5988 and, 5744 and 5739 respectively. Both PCR were performed using N16961 genomic DNA as template. These two fragments were assembled by PCR using 5738 and 5739 primers. The obtained fragment was then digested by EcoRI/PstI and cloned into EcoRI/PstI digested p7848 vector. |
| pK590 | Suicide conjugative plasmid used for deletion of *attIA* | $oriV_{R6K\gamma}$, $oriT_{RP4}$; [$Cm^R$] amplification by inverse PCR of pK584 plasmid with primers 5761 and 5762. |
| pH996 (pMP234) | Tn7 shuttle vector: pSW23T:: [Tn7R-MCS-Tn7L] | $oriV_{R6K\gamma}$, $oriT_{RP4}$; [$Cm^R$] (Val, unpublished) |
| pA684 | pUC18::$recA_{Vch}$ | *ori*ColE1; [$Carb^R$] amplification of $recA_{Vch}$ fragment by PCR using recAeco5 and recApst3 primers and N16961 genomic DNA as template. This fragment was then digested with EcoRI/PstI and cloned into EcoRI/PstI digested pUC18 (Pharmacia) vector (Krin, unpublished) |
| pK621 | pUC18::$recA_{Ec}$ | *ori*ColE1; [$Carb^R$] amplification of $recA_{Ec}$ fragment by PCR using 5779 and 5780 primers and MG1655 genomic DNA as template. This fragment was cloned by Gibson Assembly into EcoRI/PstI digested pUC18. |
| pL122 | pUC18$\Delta$*lacZα*::$recA_{Vch}$ | *ori*ColE1; [$Carb^R$] amplification by inverse PCR of the pA684 plasmid with 5799 and 5800 primers. |

| | | |
|---|---|---|
| pL123 | pUC18 *ΔlacZα::recA*$_{Ec}$ | *ori*ColE1; [Carb$^R$] amplification by reverse PCR of pA684 plasmid with VN9 and 5800 primers. |
| p7741 | pKD4 | *oriV*$_{R6K\gamma}$; [Carb$^R$, Km$^R$] (Datsenko and Wanner, 2000). |
| pL310 | pMP234*::*P$_{LAC}$-*recA*$_{EC}$-FRT-*aph*-FRT | *oriV*$_{R6K\gamma}$, *oriT*$_{RP4}$; [Cm$^R$] the P$_{LAC}$-*recA*$_{Ec}$ fragment was amplified with 5918-5916 primers from pL123 vector and the FRT-*aph*-FRT fragment was amplified with 5917-5919 primers from p7741 vector. These fragments were assembled by PCR using 5918 and 5919 primer. The obtained fragment was cloned by Gibson Assembly into pMP234 vector backbone that was amplified by inverse PCR with primers 5480 and 5481. |
| pL312 | pMP234*::*P$_{LAC}$-*recA*$_{Vch}$-FRT-*aph*-FRT | *oriV*$_{R6K\gamma}$, *oriT*$_{RP4}$; [Cm$^R$] The same cloning procedure was used as for pL310 construction. However, in this case, the P$_{LAC}$-*recA*$_{Vch}$ fragment was amplified from the pL122 vector. |
| pF324 (pMVM1) | Tn7 helper: *araC* P$_{BAD}$-*tnsABCD* | *oripSC101*ts, *oriT*$_{RP4}$; [Carb$^R$] (de Lemos Martins et al., 2018) |

*Table S3: Primers used in this study*

| Primers | Sequences |
|---|---|
| | **Primers used for Plasmid construction** |
| recAeco5 | GGAATTCGATGGACGAGAATAAACAGAA |
| recApst3 | AACTGCAGTTAAAACTCTTCTGGCACCG |
| VN9 | ATGGACGAGAATAAACAGAAGG |
| VC0542A | GGAATTCGCTTTGTGTTTGATTTCTTT |
| RECAB | CTTTGCATTCAGCCTGCCGATACTCTCTCCGGATAGTCAC |
| RECAC | GTGACTATCCGGAGAGAGTATCGGCAGGCTGAATGCAAAG |
| RECAD | GGAATTCGTGGCTGATGCCGCTTTTGA |
| 5234 | TTTCTAGGCACCAATAACTGCCTTA |

| | |
|---|---|
| 5235 | GTTGATACCGGGAAGCCCTGGGCCA |
| 5480 | GCACCTAGGAGGCGCGCCAC |
| 5481 | CGCTATTGACCCGGGATCTG |
| 5640 | GACATTATTTGCCGACTACCTTGGTGATCTCGC |
| 5641 | GATGCACTAAGCACATAATTGCTCACAGCC |
| 5738 | CGGGAATTCCCGTAGAGTAACTTGATGGGAAGTT |
| 5739 | CGGCTGCAGGAGGTCAAACTATAAAACACCCAAGC |
| 5744 | CCCTAAAACAAAGTTAGGCATCAGAGCTTTCTTGCTAATGTTAGATCAAT |
| 5761 | CTAATTAATAGACCACTGGGTGC |
| 5762 | GCTTTCTTGCTAATGTTAGATCA |
| 5779 | AGTGCCAAGCTTGCATGCCTGCAGTTAAAAATCTTCGTTAGTTTCTGC |
| 5780 | CAGCTATGACCATGATTACGAATTCGATGGCTATCGACGAAAACAAACAG |
| 5799 | ATGGCTATCGACGAAAACAAACAGAAAGC |
| 5800 | AGCTGTTTCCTGTGTGAAATTGTTATCC |
| 5914 | TAAGGCAGTTATTGGTGCCTAGAAAGTCGATGCACTAAGCACATAATTGC TCAC |
| 5915 | TGGCCCAGGGCTTCCCGGTATCAACAATTCCCACGGGTTTTGCTGCCCGC |
| 5916 | GCTCCAGCCTACACAATCGCTCAAAAACGACGGCCAGTGCCAAGCTTGC |
| 5917 | GCAAGCTTGGCACTGGCCGTCGTTTTTGAGCGATTGTGTAGGCTGGAGC |
| 5918 | CAGATCCCGGGTCAATAGCGACTGGAAAGCGGGCAGTGAGCGC |
| 5919 | GTGGCGCGCCTCCTAGGTGCATGGTCCATATGAATATCCTCC |
| 5988 | TGATGCCTAACTTTGTTTTAGGGCGACTGCCCTGCTGCGTAACATCGTTGC TGCTCCATAACATCAAACTAATTAATAGACCACTGGGTGC |
| 6176 | GGCCGAATTCCTGTGAAATCTCATGATTTCGC |
| 6177 | GGCCGGATCCGGATAGATGCATATAATTCGC |
| 6184 | TCTGTGTCGTTTTTACATCGGTATGTCC |
| 6185 | TTTTCACTCATGTTCTTGATAGAGGTGCAAGCG |
| 6178 | CCCTATGCTACTCCGTCAAGCCGTCAATTGTCTGATTCGTTACCAACACGC CTGCCAGGAATTGGGGATCGGTTAAGACCC |
| 6181 | GGGTCTTAACCGATCCCCAATTCCTGGCAGGCGTGTTGGTAACGAATCAG ACAATTGACGGCTTGACGGAGTAGCATAGGG |
| 6182 | CGCAGGGGTAGTGAATCCGCCAGGATTGACTTGCGCTGCCTTTTGCCTCCT AACTAGGTCATTTGATATGCCTCCGG |

| | |
|---|---|
| 6183 | CCGGAGGCATATCAAATGACCTAGTTAGGAGGCAAAAGGCAGCGCAAGT CAATCCTGGCGGATTCACTACCCCTGCG |
| 6229 | CCGGAGGCATATCAAATGACCTAGTTAGGAGGCAAAAATGAAATCCCAG TTTTTGTTAAGTGTTCGCGAATTTATGCAAACTCG |
| 6230 | CGAGTTTGCATAAATTCGCGAACACTTAACAAAAACTGGGATTTCATTTTT GCCTCCTAACTAGGTCATTTGATATGCCTCCGG |
| | **Primers used to identify the location of cassette insertion** |
| SWbeg | CCGTCACAGGTATTTATTCGGCG |
| SWend | CCTCACTAAAGGGAACAAAAGCTG |
| MFD | CGCCAGGGTTTTCCCAGTCAC |
| 573 | GCTGCCCGGATTACACC |
| 1366 | AGCGGGTGTTCCTTCTTCACTG |
| 1388 | CCGGGCAGGATAGGTGAAGTAG |
| 1704 | AGAGAACATAGCGTTGCCTTGG |
| 1863 | GGCCACGCGTCGACTAGTACNNNNNNNNNNACGCC |
| 1865 | GGCCACGCGTCGACTAGTAC |
| 2405 | ATACGACTCACTATAGGGCG |
| 5778 | GTCAAGAGGCTATACAGACATCAGC |
